# Mechanism of giant magnetic field effect in fluorescence of mScarlet3, a red fluorescent protein

**DOI:** 10.1101/2025.02.26.640351

**Authors:** Katherine M. Xiang, Hana Lampson, Rebecca Frank Hayward, Andrew G. York, Maria Ingaramo, Adam E. Cohen

## Abstract

Several fluorescent proteins, when expressed in *E. coli*, are sensitive to weak magnetic fields^1^. We found that mScarlet3 fluorescence in *E. coli* reversibly decreased by 21% in the presence of a 60 mT magnetic field, the largest magnetic field effect (MFE) reported in any fluorescent protein. Purified mScarlet3 did not show an MFE, but addition of flavin mononucleotide (FMN) and simultaneous illumination with blue and yellow light restored the MFE. Through extensive photophysical experiments, we developed a quantitative model of the giant MFE in mScarlet3-FMN mixtures. The key reaction step involved electron transfer from fully reduced FMNH_2_ to triplet-state mScarlet3, to form a triplet spin-correlated radical pair. The magnetic field then controlled the branching ratio between singlet recombination vs. triplet separation. Our quantitative model of the mScarlet3-FMN photocycle provides a framework for design and optimization of magnetic field-sensitive proteins, opening possibilities in fluorescent protein-based magnetometry, magnetic imaging, and magnetogenetic control.

## Introduction

Interactions between milli-Tesla magnetic fields and organic matter at room temperature are too weak to affect reaction outcomes through thermodynamic means. Nonetheless, magnetic field effects (MFEs) on photochemical reactions have long been observed in organic dye solutions.^2,3^ In linked donor-bridge-acceptor molecules, magnetic modulation of exciplex fluorescence can be as large as 80%.^4,5^ MFEs are also observed in some biochemical systems, but such effects are typically small (< 1%)^6,7^ or restricted to non-physiological conditions.^8,9^ Identification of large (>10%) MFEs in biochemical reactions under physiological conditions would open the possibility to harness these effects to modulate biochemical signals, or to implement quantum coherent measurement or control protocols in biology, and may shed light on possible mechanisms of magnetosensation.

A recent report described MFEs in several fluorescent proteins in live *E. coli*, and when paired with redox cofactors *in vitro*.^1^ Enhanced green fluorescent protein (EGFP) fused to a flavin-binding tag showed a 0.25% MFE in *E. coli* when exposed to a 25 mT field. A mixture of purified mScarlet, a red fluorescent protein, and FMN showed a nearly 3% MFE in a 10 mT magnetic field, provided that the sample had been pre-illuminated with 470 nm light. Mutagenesis and screening in a light-oxygen-voltage-sensing (LOV) domain protein led to an astonishing 35% magnetic modulation in flavin fluorescence in an engineered variant termed magLOV.^1^ The mechanisms underlying these newly discovered MFEs are not known.

MFEs are thought to arise through quantum dynamics in a spin-correlated radical pair intermediate (reviewed extensively in ^10^). Briefly, an electronic excited state — usually produced by photoexcitation — undergoes electron transfer to produce a pair of electrons that are weakly interacting but whose spin states are quantum mechanically entangled, either in a singlet or triplet state, i.e. a spin-correlated radical pair (SCRP). Each electron spin precesses under the influence of distinct hyperfine interactions with nearby nuclear spins, leading to interconversion between singlet and triplet states, known as intersystem crossing (ISC). An external magnetic field partially decouples the electron and nuclear spins, suppressing ISC. Singlet radical pairs are more likely to recombine, while triplets are more likely to separate, leading to an MFE on the photochemical reaction outcome.

While the broad outlines of the SCRP mechanism are well established, the molecular species, reaction steps, and rate constants were not known for the MFE in any fluorescent protein. An understanding of the photocycle could guide efforts to apply or enhance protein-based magnetic field effects. Ultimately, our goal is to establish a foundation for coherent control of spin dynamics in proteins.

## Results

### mScarlet3 shows a large MFE in live *E. coli*

We imaged the fluorescence of several red fluorescent proteins expressed in live *E. coli* using an epifluorescence microscope. We used a servo-controlled magnet on a stick to switch the magnetic field between 0 and 60 mT (Fig. 1A). We tested red-emitting fluorescent proteins mScarlet3, mScarlet-I3, mSandy2, mRuby3, and mKate2. All except mKate2 showed magnetically modulated fluorescence (Fig. S1).

**Figure 1:**
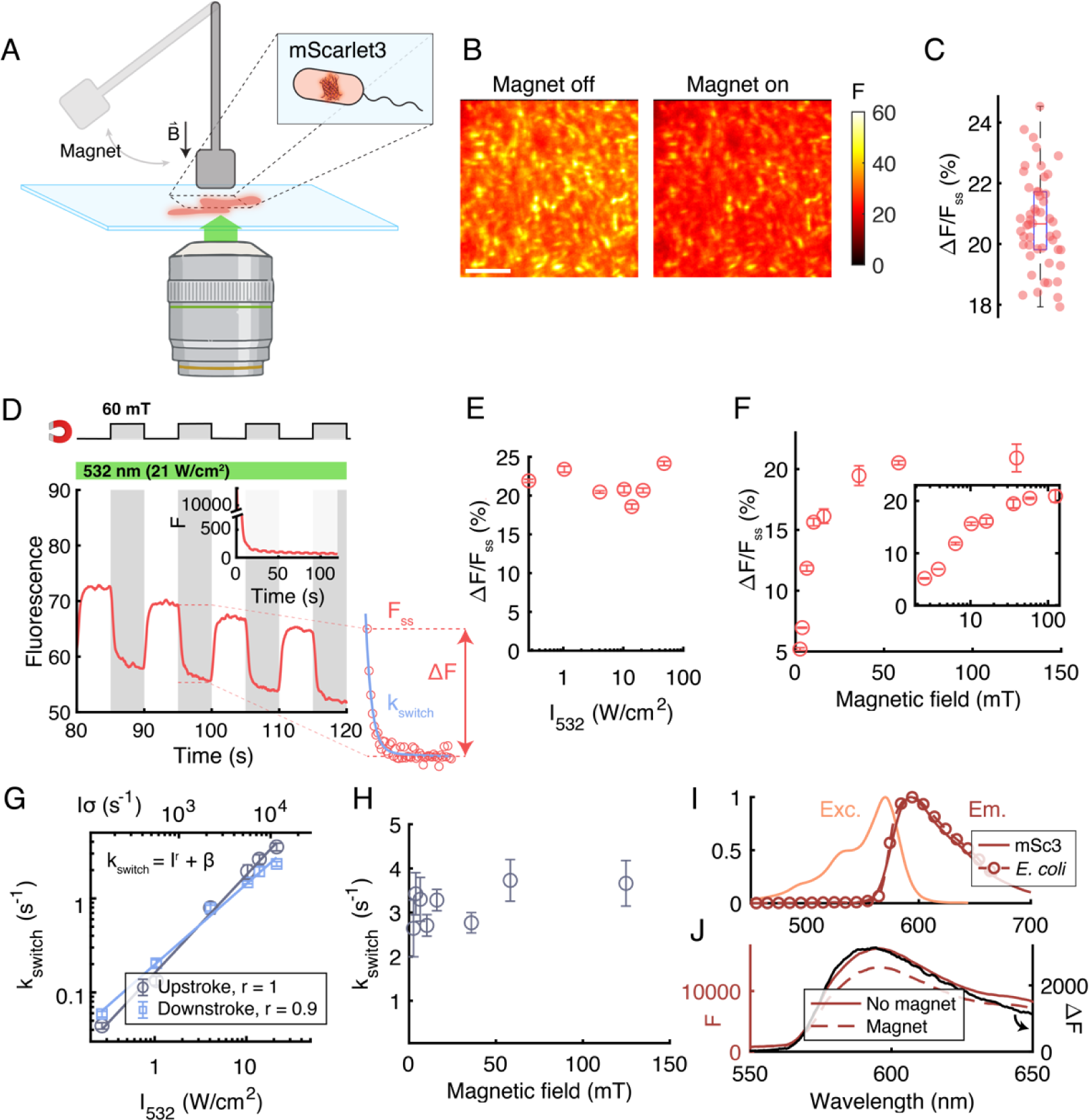
mScarlet3 fluorescence in E. coli is magnetic field sensitive. A) Experimental setup. Fluorescence of *E. coli* expressing mScarlet3 was probed via epifluorescence microscopy, while a magnet on a servo modulated the local magnetic field. B) Image of *E. coli* expressing mScarlet3 in the absence (left) or presence (right) of a 60 mT magnetic field. Same contrast for both images. Scale bar 10 µm. C) Distribution of amplitudes of the MFE across individual *E. coli*. ΔF/F_ss_ = (20.7 ± 1.5)% (mean ± s.d., *n* = 50 cells). Boxplot shows median, 25^th^ and 75^th^ percentiles; whiskers show extrema. D) Fluorescence time-trace of mScarlet3 under green excitation (532 nm, 21 W/cm^2^) and alternating magnetic field. Inset: Upon illumination onset, fluorescence rapidly quenched. Right: definitions of the parameters *k*_switch_, ΔF, and F_ss_. E) Steady state MFE amplitude as a function of illumination intensity. Magnetic field was 60 mT. F) Steady state MFE amplitude as a function of magnetic field. Illumination intensity was 21 W/cm^2^. G) Switching rate as a function of illumination intensity. Magnetic field was 60 mT. Rates were approximately first-order in illumination intensity for both the upstroke and the downstroke. H) MFE switching rate for the upstroke as a function of magnetic field. Illumination intensity was 21 W/cm^2^. Rates were independent of magnetic field. I) Excitation and emission spectra of purified mScarlet3 (solid curve), overlaid with emission spectrum of *E. coli* expressing mScarlet3 (circles). J) The magnetic field modulated the amplitude (red traces), but not the shape (black trace), of the fluorescence emission spectrum. Error bars for E–H denote s.e.m.

We focused on mScarlet3 because it was bright and had the largest MFE of the proteins we tested. In the presence of the magnetic field (*B* = 60 mT), the fluorescence of mScarlet3 was visibly dimmer than in its absence (Fig. 1B; 532 nm excitation, 48 W/cm^2^, 575 nm long-pass emission, *B* switching every 5 s). We quantified the MFE amplitude by ΔF/F_ss_, where F_ss_ is the steady-state fluorescence at magnetic field *B* = 0, and ΔF = F_ss_ – F(*B*). Quantification across individual mScarlet3-expressing bacteria showed an MFE of ΔF/F_ss_ = 20.7 ± 1.5% (mean ± s.d., *n* = 50 cells, Fig. 1C).

We then explored how the illumination intensity and the magnetic field strength affected the dynamics of the fluorescence. Upon onset of illumination at 21 W/cm^2^, the fluorescence rapidly quenched from 13,000 to 80 counts, with a time-constant of ∼1.8 s (Fig. 1D inset). After the fluorescence stabilized, step-wise changes in magnetic field induced gradual changes in fluorescence. The steady-state MFE amplitude was independent of the excitation intensity over the measured range *I* = 0.3 – 48 W/cm^2^ (Fig. 1E, Fig. S2). The steady-state MFE amplitude increased with increasing magnetic field, with a half-saturation field *B*_1/2_ = 5.5 mT (Fig. 1F).

We fit the magnetic field-induced switching rate constant, *k_switch_* (defined in Fig. 1D), over an intensity range from *I* = 0.3 – 21 W/cm^2^ (corresponding to per-molecule excitation rates *Iσ* = 145 to 11,600 s^-1^; *σ* is the absorption cross-section at 532 nm and is proportional to *ε*_532_ = 53,040 M^-1^cm^-1^). The switching rate at magnetic field onset varied from 0.04 to 3.5 s^-1^ over this range, and was approximately linear in intensity, i.e. *k*_switch_ ∝ *I*^*r*^ + *β* with *r* = 1.0 for the fluorescence upstroke and *r* = 0.9 for the downstroke (Fig. 1G). These results suggested a single rate-limiting photochemical step in the magnetic-field dependent photochemical reaction. The ratio of k_switch_ (s^-1^) to per-molecule excitation rate *Iσ* (s^-1^) gives a branching ratio of 3.1 × 10^-4^ for magnetic field-dependent photoswitching from bright to dark forms of mScarlet3.

The switching rate was independent of the magnetic field strength over the range 3 – 125 mT (Fig. 1H), implying that the magnetically modulated rate constant(s) were not the rate-limiting steps in the photochemical conversion between fluorescent and non-fluorescent mScarlet3 states. The fluorescence emission spectrum of the *E. coli* closely matched the spectrum for purified mScarlet3 (Fig. 1I),^11^ and the MFE modulated the amplitude, but not the shape, of this spectrum (Fig. 1J). These observations indicate that mScarlet3 was the primary emissive species.

In *E. coli*, various factors could influence the MFE including pH, redox factors, and diffusional confinement of molecules within each bacterium. We found that the MFE was largest on the second day of expression in liquid culture, suggesting the involvement of metabolic or redox factors beyond just the fluorescent protein. To study the molecular mechanisms underlying the MFE we thus switched to working with purified protein.

### Purified mScarlet3 shows an MFE in the presence of FMN and blue light

We purified mScarlet3 and measured its fluorescence while varying the magnetic field. An imaging spectrometer recorded both the amplitude and shape of the emission spectrum (Fig. 2A). No MFE was detected for yellow (561 nm) or yellow + blue (561 nm + 488 nm) excitation (Fig. 2B). Prior work showed that the addition of flavin mononucleotide (FMN, Fig. 2C) and blue light restored the MFE in mScarlet3,^1^ so we replicated that result. Addition of 60 µM FMN did not produce an MFE under yellow light alone. However, when we combined blue and yellow light in the presence of FMN, we observed a rapid initial drop in mScarlet3 fluorescence, and then an MFE (Fig. 2D). The blue light also evoked FMN fluorescence, which was spectrally distinct from mScarlet3 fluorescence. The FMN fluorescence showed a rapid initial drop before stabilizing, but did not show a detectable MFE. Upon turning off the blue light, the mScarlet3 fluorescence partially recovered, and the MFE went away.

**Figure 2.**
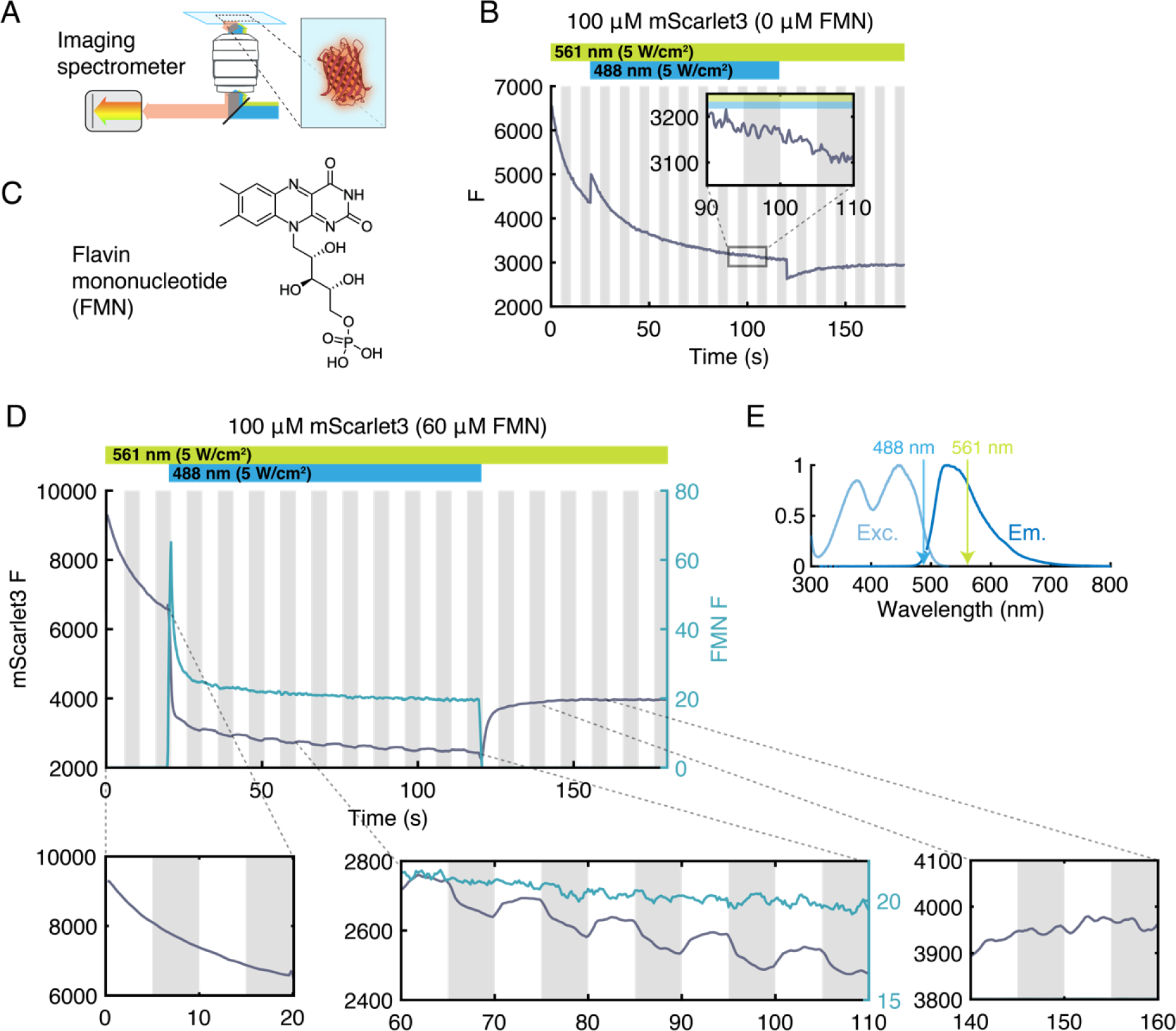
The mScarlet3 MFE requires FMN and blue light. A) Fluorescence spectra were measured in a microscope with an imaging spectrometer. B) Fluorescence of purified mScarlet3 showed no MFE under yellow (561 nm) or yellow + blue (488 nm) illumination. The magnetic field was modulated in 5 s intervals between *B* = 60 mT (grey stripes) and *B* = 0 (white stripes). C) Chemical structure of FMN. D) Addition of FMN (60 µM) did not affect mScarlet fluorescence under yellow-only illumination. Onset of blue illumination led to partial quenching of mScarlet3 fluorescence and appearance of an MFE. Fluorescence was spectrally filtered to collect mScarlet3 emission and to reject flavin emission. E) Excitation and fluorescence emission spectra of FMN.

Based on these measurements, and the overlap of the blue laser wavelength with the FMN absorption spectrum (Fig. 2E), we speculated that a photochemical product of FMN excitation might be involved in the MFE. To test this hypothesis, we prepared a sample comprising mScarlet3 immobilized on Ni-NTA resin beads, immersed in a solution of FMN (500 µM), and sandwiched between glass coverslips. We targeted the yellow light to focus on a single bead, and the blue light to pass through a region of FMN solution approximately 200 µm from the bead (Fig. 3A,B). This arrangement prevented any direct blue light interaction with mScarlet3 molecules, while allowing diffusive interchange of long-lived photoproducts between the two illuminated regions. We measured the emission from both the blue-illuminated and yellow-illuminated regions with an imaging spectrometer (Fig. 3B, right), while we varied the magnetic field.

**Figure 3.**
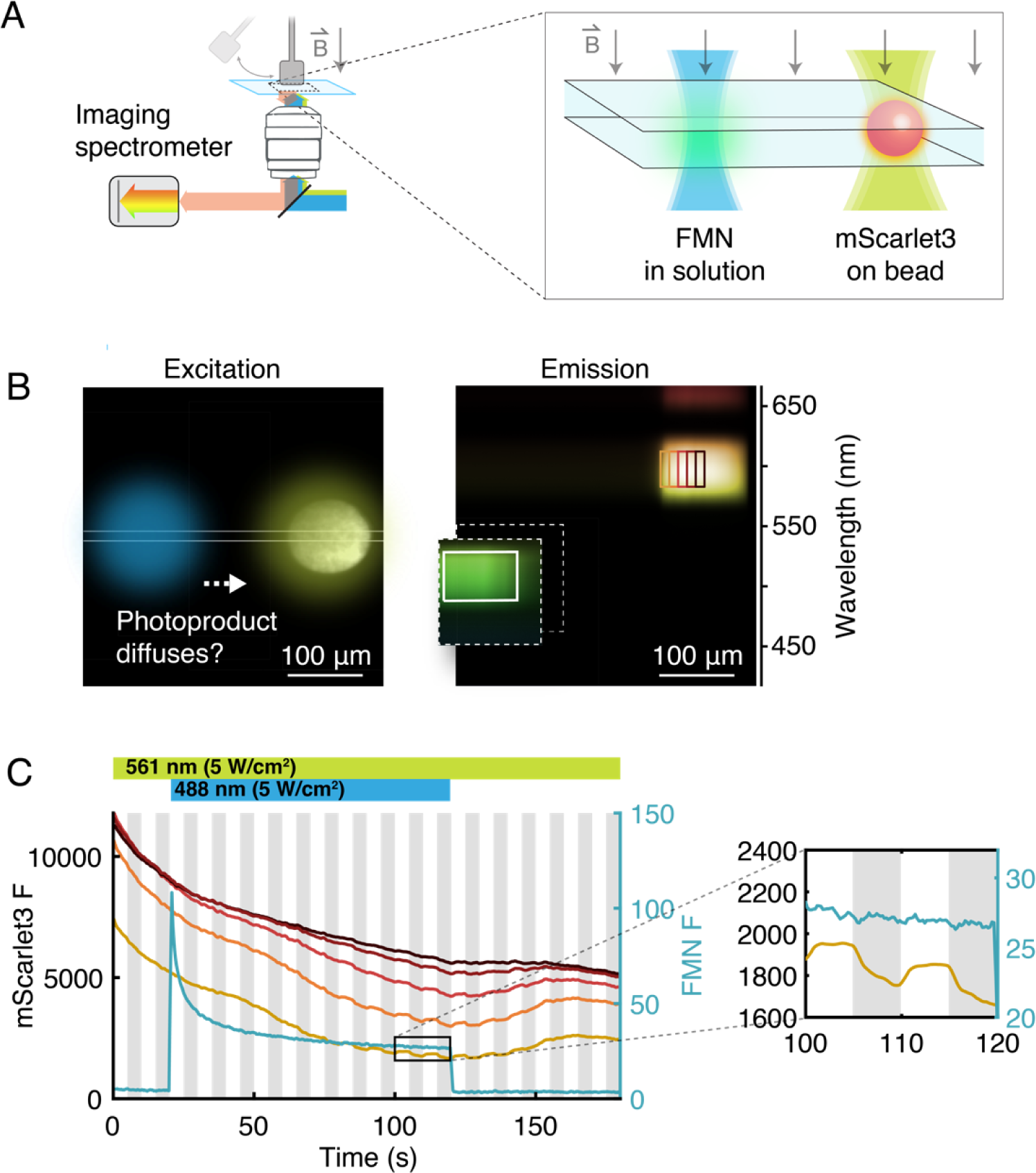
The mScarlet3 MFE requires a long-lived FMN photoproduct. A) Experiment to test whether an FMN photoproduct contributed to the MFE. A bead containing tethered mScarlet3 was illuminated with yellow light. Blue light was laterally displaced and focused into a solution of FMN. B) Left: Excitation profiles overlaid on an image of the bead. Right: Emission of FMN (green) and mScarlet3 (yellow) on an imaging spectrometer. The x-axis corresponds to position and the y-axis to emission wavelength. The contrast of the FMN emission has been increased to render it visible. C) Time-course of FMN fluorescence (cyan) and mScarlet3 fluorescence in the regions on the bead indicated with correspondingly colored rectangles in (B). The magnetic field was modulated in 5 s intervals between *B* = 60 mT (grey stripes) and *B* = 0. Approximately 35 s after blue light onset, the mScarlet3 fluorescence showed quenching and onset of an MFE, with the regions closer to the blue light responding first. The FMN fluorescence was not magnetic field sensitive.

As with the soluble proteins, the mScarlet3 initially showed no MFE under yellow-only illumination. When the blue light was turned on, the FMN fluorescence decayed over ∼15 s, suggesting photochemical consumption of the FMN. As before, there was no MFE in the FMN fluorescence. After a delay of ∼35 s, the mScarlet3 fluorescence began to drop, and an MFE appeared (Fig. 3C). The side of the bead closest to the blue spot had the earliest and largest response, and more distant regions had later and smaller responses. When the blue light was turned off, the MFE persisted for ∼35 s, and then gradually faded as the mScarlet3 fluorescence recovered.

These experiments taught us several important facts. First, they established that a relatively long-lived FMN photoproduct was necessary for the MFE. Second, the magnet did not act on the blue-mediated FMN photochemistry, but rather on the interaction of the FMN photoproduct with yellow-excited mScarlet3. Third, the MFE did not require blue light interaction with mScarlet3 itself (or with mScarlet3 photoproducts). We estimated the diffusion coefficient of the FMN photoproduct from *D* ≈ *x*^2^/2*t*, where *x* = 200 µm was the distance between the blue laser spot and the mScarlet3-labeled bead, and *t* = 35 s was the delay between blue illumination and onset of the MFE. This estimate gave *D* ≈ 570 µm^2^/s, broadly consistent with diffusion of a small molecule in water (due to the complex geometry, this estimate for *D* is likely only accurate within a factor of ∼2).

### The mScarlet3 MFE requires both oxidized and reduced FMN

Optically excited FMN is a potent oxidizing agent, readily forming the semiquinone radical. This radical spontaneously disproportionates to re-form FMN and the doubly reduced FMNH_2_ (Fig. 4A).^12–16^ We speculated that photochemically produced FMNH_2_ might mediate the MFE.

**Figure 4.**
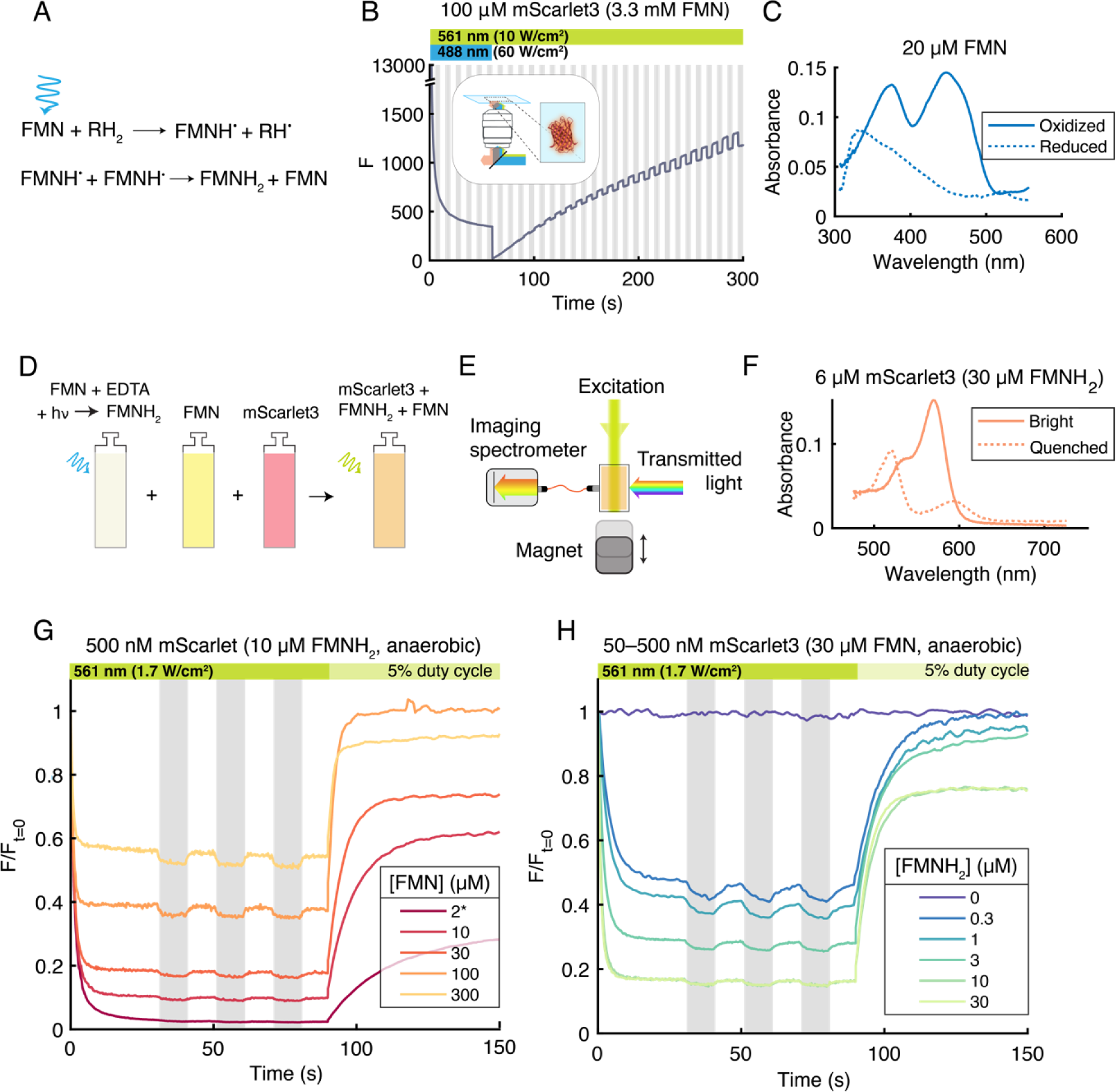
The mScarlet3 MFE requires both oxidized and reduced FMN. A) Photo-excitation of FMN produces FMNH_2_. B) Fluorescence of purified mScarlet3 with an excess of FMN and high-intensity blue light. The fluorescence quenched almost completely (residual fluorescence during blue illumination is the tail of the FMN emission and autofluorescence due to the high blue intensity). Upon turning off the blue light, the mScarlet3 fluorescence recovered and the MFE increased proportionally. C) Absorption spectra of FMN in its oxidized and photoreduced (FMNH_2_) states. D) Samples of FMNH_2_, FMN, and mScarlet3 were prepared separately in anaerobic conditions, and then combined and measured under 561 nm illumination. E) Top view of the experimental apparatus. Magnetic field-dependent fluorescence and absorption spectra were recorded on a spectrometer. F) Absorption spectra of mScarlet3 and the fully photoreduced mScarlet3 product (presumably mScarlet3^•-^ radical anion) in the presence of FMNH_2_. G, H) mScarlet3 fluorescence quenching, MFE, and recovery in samples with indicated concentrations of FMN and FMNH_2_. All fluorescence traces are normalized to their initial value. A 5% duty cycle of yellow light (25 ms every 0.5 s) was used to image the fluorescence recovery. Fluorescence during this epoch was scaled 20× to account for the lower illumination intensity. Asterisk indicates FMN concentration determined from absorption measurements (Methods, Fig. S3).

To test this hypothesis, we produced FMNH_2_ by illuminating FMN (3.3 mM) with high intensity 488 nm light (60 W/cm^2^), in a solution also containing mScarlet3 (100 µM). As before, we used yellow light (561 nm, 10 W/cm^2^) to probe the mScarlet3 fluorescence. To our surprise, the mScarlet3 fluorescence rapidly quenched to almost zero, and the MFE was initially not detectable (Fig. 4B). This observation seemed to contradict our hypothesis. However, after the blue light was turned off, the fluorescence gradually increased, and the MFE appeared, growing proportionally with the fluorescence. We hypothesized that ambient oxygen was re-oxidizing the FMNH_2_ to FMN, and that perhaps it was necessary to have both FMNH_2_ and FMN in the solution to observe the MFE.

To test this revised hypothesis, we produced FMNH_2_ by illuminating FMN with 488 nm light in the presence of 5 mM ethylenediaminetetraacetic acid (EDTA), an efficient electron donor for photoreduction of FMN^14,16,17^. We performed the photoreduction in an anaerobic glovebox to avoid reoxidation. Absorption spectra confirmed photoreduction of FMN to FMNH_2_ (Fig. 4C). We then prepared mixtures with degassed solutions containing well-defined concentrations of FMNH_2_, FMN, and mScarlet3 under anaerobic conditions (Fig. 4D), and sealed the samples for characterization.

We used a home-built apparatus to interleave measurements of the absorption and fluorescence emission spectra while exciting the mScarlet3 with yellow light and varying the magnetic field (Fig. 4E). In a sample of 30 µM FMNH_2_ and 6 µM mScarlet3 (and no FMN), illumination with 561 nm light (1.7 W/cm^2^) caused a rapid quenching of the mScarlet3 fluorescence to almost zero. The absorption spectrum of the mScarlet3 photoproduct was distinct from unquenched mScarlet3 (Fig. 4F) with both a blue-shifted and red-shifted peak. We ascribe this spectrum to a photoreduction of mScarlet3 by FMNH_2_, most likely to form the mScarlet3^•-^ radical anion. The putative mScarlet3^•-^ radical anion was not fluorescent under 561 nm excitation.

We then systematically varied the concentrations of FMN and FMNH_2_ in mScarlet3 solutions and recorded the magnetic field-dependent fluorescence (Fig. 4G–H). When [FMNH_2_] was 1 µM or less, we used 50 nM mScarlet3 to ensure that FMNH_2_ was in excess; otherwise, we used 500 nM mScarlet3. Complete photoreduction from FMN to FMNH_2_ is difficult^19^, so for the samples with low [FMN] we determined the concentrations of each flavin species from absorption spectra (Fig. S3). In contrast to previous experiments, the MFE only required yellow light.

Increasing [FMNH_2_] decreased steady-state mScarlet3 fluorescence, and increasing [FMN] counteracted this effect. The MFE was largest in absolute magnitude when these two effects were balanced so the steady-state fluorescence was approximately half its initial value. At fixed [FMNH_2_], increasing [FMN] increased the rate of initial quenching and also the rate of magnetic field-induced switching (Fig. 4G). In the absence of FMNH_2_, FMN had no effect on mScarlet3 and did not induce an MFE (Fig. 4H, purple trace), as we observed previously (e.g. Fig. 2B).

After 90 s of yellow illumination, we decreased the yellow intensity to 5% (85 mW/cm^2^), to monitor the recovery of mScarlet3 fluorescence in close-to-dark conditions. The rate of mScarlet3 recovery in dim yellow light depended on [FMN] and was largely independent of [FMNH_2_] (except for the small effect of residual yellow-light + FMNH_2_-mediated quenching).

In anaerobic conditions, the MFE persisted overnight (Fig. S4A), establishing that the active FMN photoproduct was stable in the absence of oxygen. All evidence was consistent with FMNH_2_ as the photoproduct responsible for photo-induced mScarlet3 quenching and the MFE. To rule out the possibility of species other than FMNH_2_ and FMN contributing to the MFE, we tested mixtures of mScarlet3 with EDTA alone (Fig. S4B), as well as with FMN photoproducts lumichrome and H_2_O_2_ (Fig. S4C).^15–17^ None of these mixtures showed an MFE. We measured the MFE at several pH values to determine whether the reductant was FMNH_2_ or FMNH^-^, as FMNH_2_ has a pK_a_ of 6.7^18^. The mScarlet3 fluorescence quenched more at lower pH, suggesting that FMNH_2_ was the relevant species (Fig. S5).

Together, these results imply that FMNH_2_ electrochemically reduced the optically excited mScarlet3 to form a non-fluorescent protein photoproduct, and FMN re-oxidized this photoproduct to re-form ground-state mScarlet3. In this model, electron transfer from FMNH_2_ to mScarlet3 likely proceeded through a radical-pair intermediate, whose branching between recombination and separation was magnetic field sensitive.

### The mScarlet3 photoreduction likely proceeds through an mScarlet3 triplet state

Two observations led us to suspect that FMNH_2_ electrochemically reduced photogenerated mScarlet3 triplet (^3^mSc) rather than the excited mScarlet3 singlet (mSc*). First, under our illumination conditions, the population of mSc* was very low. The mSc* lifetime is 4 ns^11^, and at an excitation intensity of 1.7 W/cm^2^ at 561 nm, the per-molecule excitation rate was *Iσ* = 1652 s^-1^ (*ε*_561_ = 90,480 M^-1^cm^-1^). Thus, the excited-state to ground-state ratio mSc*/mSc ≈ 6 × 10^-6^. At 10 µM FMNH_2_, and a measured rate of initial quenching of 0.98 s^-1^, the implied bimolecular rate constant if the reaction proceeded from mSc* would be 8.3 × 10^10^ M^-1^ s^-1^, substantially greater than the diffusion-limited aqueous bimolecular rate constant, 6.6 × 10^9^ M^-1^ s^-1^.^19^ Hence a reaction from a long-lived triplet which could accumulate to higher concentration was more likely.

Second, we noted that the magnet decreased the mScarlet3 fluorescence. A magnetic field prolongs the lifetime of the initial spin state of a SCRP. A singlet SCRP is more likely to recombine, releasing ground-state mScarlet3, which can fluoresce, while a triplet SCRP is more likely to diffusively separate, building up a population of non-fluorescent mSc^•-^ radical anion. Thus we surmised that the SCRP was likely born in a triplet state. The electron-transfer reaction usually conserves spin, so one of the reactants was likely a triplet. FMNH_2_ has a singlet ground state, so the mScarlet3 was likely in a triplet state.

We measured the mSc* ➔ ^3^mSc intersystem crossing and ^3^mSc ➔ mSc recovery rates for mScarlet3 (with no flavin or other redox factors present), which we modeled as a three-state system (Fig. 5A). In an anaerobic sample of mScarlet3 bound to beads (to eliminate effects of diffusion), we applied pulses of 561 nm light at different intensities (Fig. 5B) and recorded the millisecond-timescale fluorescence dynamics (Fig. 5C). We observed a reversible intensity-dependent initial dip in the fluorescence, which we attributed to populating the mScarlet3 triplet state. We quantified the rate (Fig. 5D) and amplitude (Fig. 5E) for this initial dip as a function of the mScarlet3 excitation rate (*Iσ*), where *I* is the excitation intensity (photons/cm^2^/s) and *σ* is the absorption cross section at 561 nm (cm^2^). From these, we determined 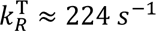 and *k*_ISC_ ≈ 1.9 × 10^5^ s^−1^ (Supplementary Information).

**Figure 5.**
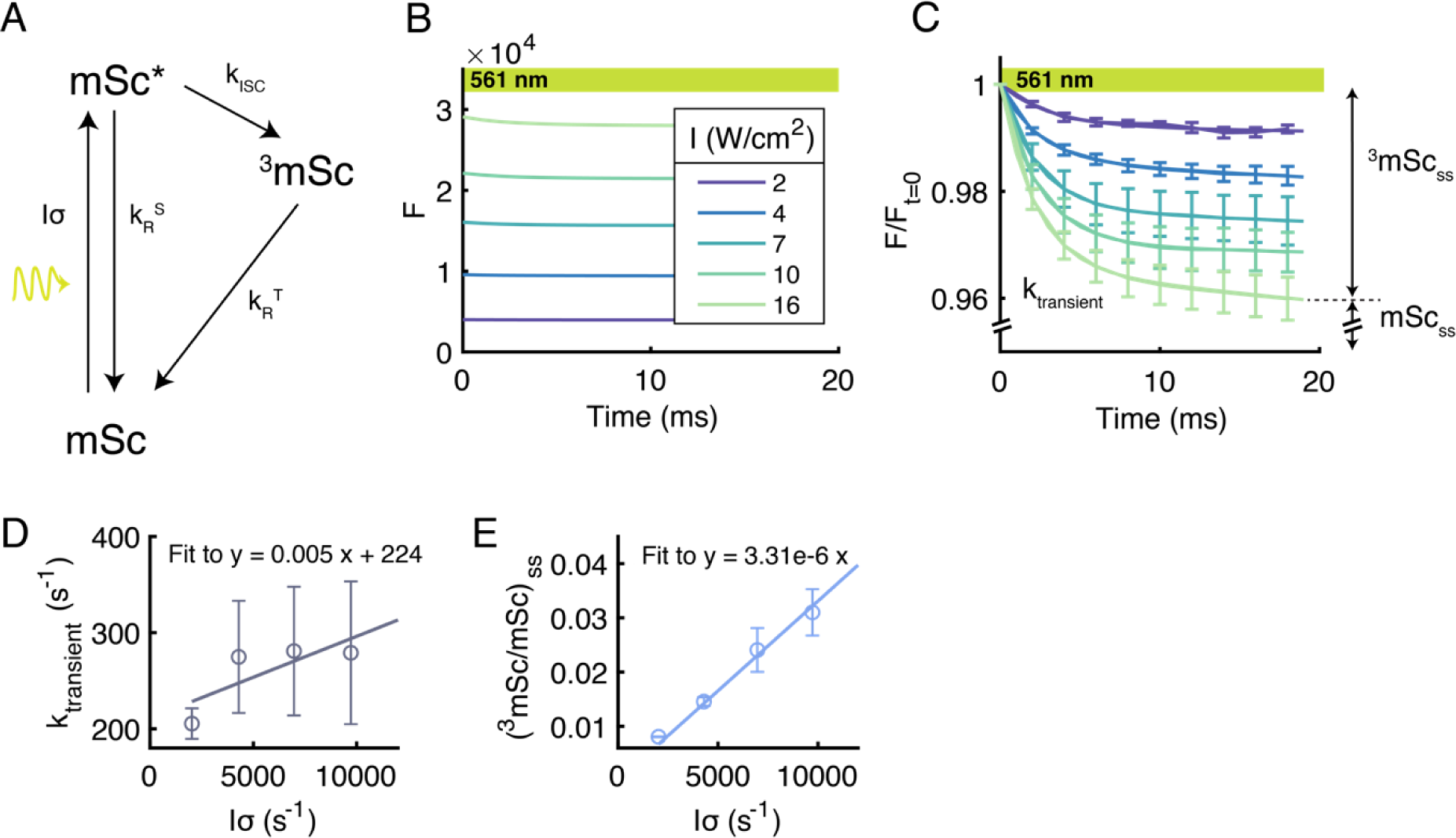
Illumination of mScarlet3 populates a transient dark state. A) Three-state model of intersystem crossing (ISC) in mScarlet3. B) Onset of high-intensity illumination of mScarlet3 immobilized on an agarose Ni-NTA bead induced a rapid and reversible slight decrease in fluorescence. C) Data from (B), normalized to initial fluorescence, with exponential + linear fits. More intense illumination led to a larger fractional decrease in fluorescence. The relative amplitudes of the fluorescence dip and the fluorescence steady-state correspond to the relative triplet and ground-state populations (assuming negligible singlet excited state population). Traces are averaged over 18 flashes for each of *n* = 3 beads. Fitting the D) rate constant and E) amplitude of the fluorescence dip implied *k*_R_^T^ ≈ 224 s^-1^ and *k*_ISC_ ≈ 1.9 × 10^5^ s^-1^. Error bars denote s.e.m.

### Quantitative model of the mScarlet3/FMNH_2_/FMN magnetic field effect

Combining all the above results, we developed a model of the mScarlet3 and FMN magnetic field effect (Fig. 6A). In a solution initially containing only FMN, the blue illumination produced FMNH_2_. Alternatively, the FMNH_2_ can be produced separately and added to the solution under anaerobic conditions. Yellow excitation populated the mScarlet3 triplet state (^3^mSc, Fig. 5). In the key reaction step, FMNH_2_ donated an electron to the mScarlet3 triplet state, producing an initially triplet spin-correlated radical pair, ^3^[mSc^•-^+ FMNH^•^]. The magnetic field suppressed intersystem crossing to the singlet. The radical pair either recombined (more likely if in a singlet) or separated (more likely if in a triplet). Once separated, FMN re-oxidized mSc^•-^ back to mScarlet3. This model contrasts with the MFE in interactions of FMN with a non-fluorescent protein backbone, where the optically excited FMN is proposed to accept an electron from the protein^7^.

**Figure 6.**
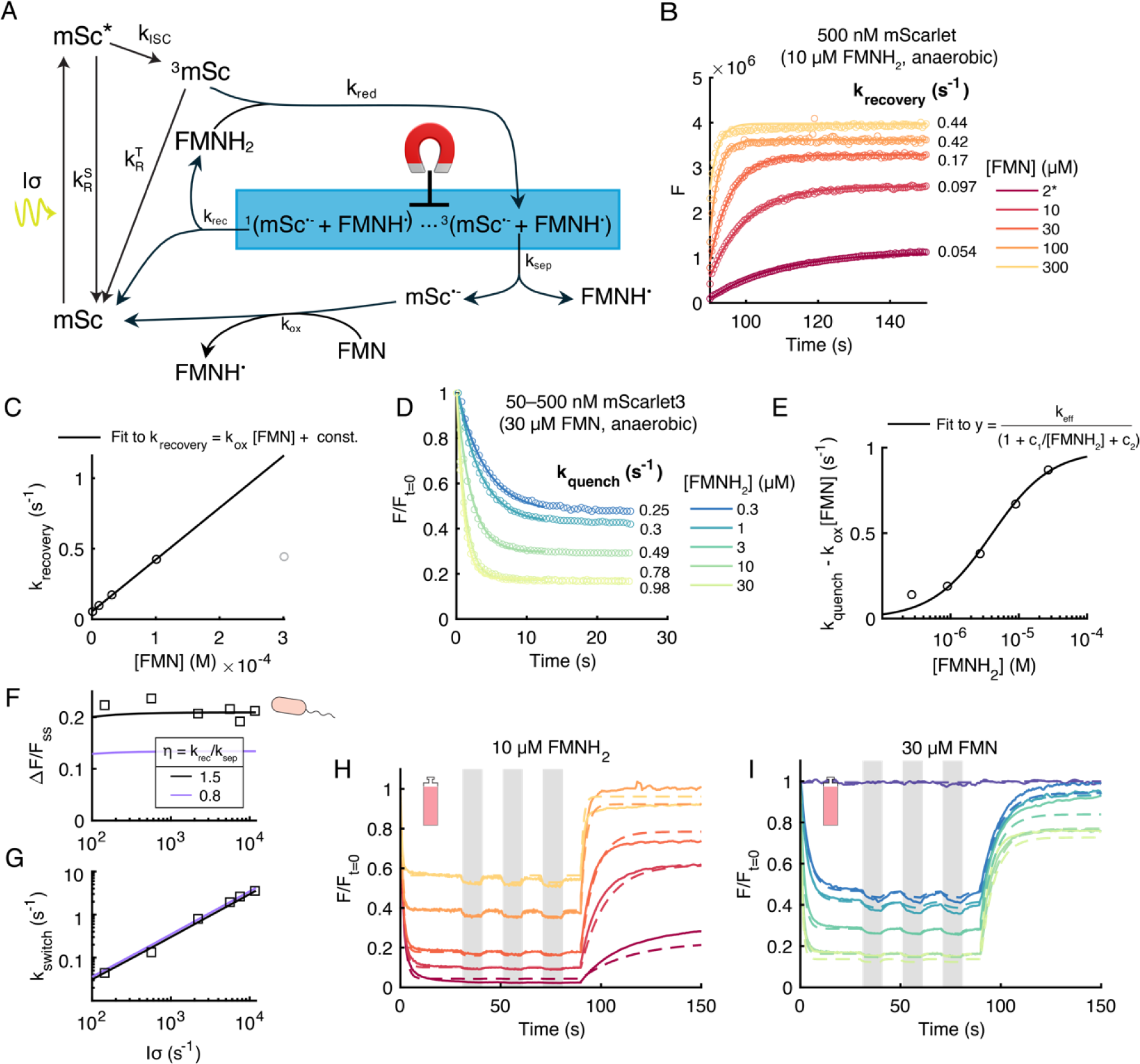
Quantitative model of giant MFE in mixture of mScarlet3, FMN, and FMNH_2_. A) Proposed mechanism. A yellow photon excites mScarlet3. The excited singlet state either fluoresces or undergoes ISC to produce the mScarlet3 triplet. Blue light drives photoreduction of FMN to FMNH_2_ (reactions shown in Fig. 4A). FMNH_2_ donates an electron to triplet mScarlet3 to form a triplet-born spin-correlated radical pair (SCRP). The magnetic field controls the rate of ISC in the SCRP. Singlet products recombine to re-form ground-state mScarlet3. Triplet products diffuse apart to release a non-fluorescent mScarlet3^•-^ radical anion. FMN re-oxidizes the mScarlet3^•-^ back to the mScarlet3 ground state. B) Exponential + constant fits of the fluorescence recovery for the FMN titration in Fig. 4G. C) Linear fit of *k*_recovery_ as a function of [FMN] determines the rate constant k_ox_ of mScarlet3^•-^ reoxidation by FMN. D) Exponential + constant fits of the fluorescence quenching for the FMNH_2_ titration in Fig. 4H. E) Fitting *k*_quench_ as a function of [FMNH_2_] determines parameters *c_1_* and *c_2_* (Supplementary Information) which yield the effective rates of the SCRP separation and recombination. F) Comparison of measured and predicted MFE amplitude as a function of light intensity in *E. coli*. G) Comparison of measured and predicted MFE upstroke switching rate as a function of light intensity in *E. coli*. H, I) Comparison of measured and predicted fluorescence dynamics for the experiments in Figs. 4G, H. All rate constants in (F) – (I) were identical. *ƞ* ≡ k_rec_/k_sep_ was fit separately for *E. coli* and *in vitro* data.

We wrote the rate equations for this model and determined the rate constants from our data. The key steps are outlined below, the details are in the Supplementary Information. We fit the rates of fluorescence recovery under dim yellow light, *k*_recovery_, in the FMN titration data (Fig. 6B). From this, we calculated the bimolecular rate constant, *k*_ox_, of mSc^•-^ reoxidation by FMN (Fig. 6C). We then fit the rates of initial fluorescence quenching, *k*_quench_, from the FMNH_2_ titration data (Fig. 6D). By combining *k*_ISC_, determined above, with the dependence of *k*_quench_ on [FMNH_2_], we determined 1) the bimolecular rate constant *k*_red_, for ^3^mSc reduction by FMNH_2_ (Fig. 6E), and 2) a constraint on the zero-field branching ratio between SCRP recombination vs. separation pathways (Supplementary Information Section 4.2).

The combined spin and reaction dynamics in the SCRP are described by the Haberkorn master equation formalism^20^. We made the simplifying approximation that the SCRP reached a quasi-steady-state S:T ratio on a short timescale compared to either of the subsequent S or T reactions. We defined *α*(B) and 1 − *α* (B) as the quasi-steady-state SCRP triplet and singlet fractions, respectively, where *B* is the magnetic field strength. Within this model, there is an indeterminacy between *α* and the ratio *ƞ* = *k*_rec_/*k*_sep_, where *k*_rec_ and *k*_sep_ are the rate constants for subsequent recombination and separation, respectively.

Theoretical predictions based on Gaussian random hyperfine fields give quasi steady-state ratios for a T_0_-born radical S:T = 2/9:7/9 at B = 0, and S:T = 1/6:5/6 in saturating magnetic field^21^. We fixed *α*(0) = 7/9 in accord with this prediction. The fit in Fig. 6E then yielded *ƞ* = 0.84 and *k_red_* = 5.1 × 10^7^ M^-1^ s^-1^. We let *α*(B_sat_) be a fitting parameter, which we constrained by fitting the numerical model to our data on ΔF/F_ss_ and ΔF/F_t=0_ as a function of [FMN] and [FMNH_2_] (i.e. Figs. 4H, I). The best-fit value of *α*(B_sat_) was 0.92 (vs. the theoretical 5/6 = 0.833). The fit parameters are given in Table 1 and the model is available as Supplementary Code.

**Table 1:** Fit parameters for the kinetic model.

| Parameter | Value | Description | Source |
| --- | --- | --- | --- |
| $\sigma$ | $3.5 \times 10^{-16} \text{ cm}^2$ | mScarlet3 absorption cross section at 561 nm | Ref <sup>11</sup> |
| $k_R^S$ | $2.5 \times 10^8 \text{ s}^{-1}$ | Inverse of the mScarlet3 excited-state lifetime | Ref <sup>11</sup> |
| $k_{ISC}$ | $1.85 \times 10^5 \text{ s}^{-1}$ | mScarlet3 intersystem crossing rate | Fit from Fig. 5E |
| $k_R^T$ | $224 \text{ s}^{-1}$ | Inverse of the mScarlet3 triplet lifetime | Fit from Fig. 5D |
| $k_{ox}$ | $3375 \text{ M}^{-1} \text{ s}^{-1}$ | Oxidation of mScarlet3 radical anion by FMN | Fit from Fig. 6C |
| $\alpha$ | $7/9 (B = 0)$<br>$0.92 (B = B_{sat})$ | Triplet fraction in the SCRP (quasi-steady-state) | Ref <sup>21</sup> (for $B = 0$);<br>Fit to numerical model for $B = B_{sat}$ |
| $k_{red}$ | $5 \times 10^7 \text{ M}^{-1} \text{ s}^{-1}$ | Reduction of triplet mScarlet3 by FMNH<sub>2</sub> | Fit from Fig. 6E |
| $\eta$ | 0.84 in solution<br>1.5 in *E. coli* | Ratio of SCRP recombination to separation | Fit from Fig. 6E (in solution) |
| $k_{dis}$ | $5 \times 10^9 \text{ M}^{-1} \text{ s}^{-1}$ | Rate constant for FMNH• disproportionation | Ref <sup>13</sup> |

Our model explained many features of our data, both in *E. coli* and with purified protein. In *E. coli*, upon onset of illumination at *Iσ* = 11,400 s^-1^, the mScarlet3 fluorescence initially dropped by more than 100-fold, from 13,000 to 80 counts. Our model reproduced the near-independence of MFE amplitude (ΔF/F_ss_) on illumination intensity under conditions of strong quenching (Fig. 6F, compare to Fig. 1E), the linear dependence of magnetic field-induced switching rate (*k*_switch_) on illumination intensity (Fig. 6G, compare to Fig. 1G), and the independence of *k*_switch_ on magnetic field strength (Fig. S6, compare to Fig. 1H). The dependence of MFE upstroke switching rate on light intensity in *E. coli* implied [FMNH_2_] ≈ 6 µM, though it is possible other redox factors contributed. The residual fluorescence after quenching implied [FMN] ≈ 0.4 µM, consistent with literature values^22^. Using the rate constants determined from our experiments in purified proteins, the maximal predicted MFE was 14% across all illumination intensities and concentrations of FMN and FMNH_2_. To match the larger MFE (21%) observed in *E. coli*, we increased the ratio of *k_*r*ec_* /*k*_sep_ from *ƞ* = 0.8 to *ƞ* = 1.5. This modification is plausible, considering the influence of nanoscale confinement and crowding in the *E. coli* cytoplasm.

For the experiments with purified protein, the model reproduced the dependence of quenching amplitude (F_ss_/F_t=0_) on FMN and FMNH_2_ concentrations (Fig. S7). We then simulated the complete experiment of Figs. 4F, G, including the initial quenching, the magnetic field-dependent changes in fluorescence, and the recovery of fluorescence under dim yellow light. By varying the nominal flavin concentrations by < 30% (to account for possible background redox processes) we achieved close overlay of the predicted and measured traces (Fig. 6H, I).

### Implications of the model

We used the kinetic model to explore how to maximize the MFE amplitude and kinetics. The most easily controlled parameters are [FMN], [FMNH_2_], and light intensity (*Iσ*). Surprisingly, we found that variation in all three parameters caused the absolute MFE, Δ*F*/*F*_*t*=0_, to vary along a single universal curve parameterized by *x* ≡ *F*_*ss*_/*F*_*t*=0_ i.e. the steady-state residual fluorescence. Specifically, in the limit of small Δα ≡ *α*(*B*_sat_) – *α*(0), the kinetic equations predicted that Δ*F*/*F*_*t*=0_ ∝ *x*(1 − *x*). *In vitro* data with variable [FMN], [FMNH_2_] (Fig. 4G, H) and *Iσ* (Fig. S8) all fell close to this curve, confirming its universality (Fig. 7A). This result confirms our earlier observation that the absolute MFE was maximized around 50% quenching.

**Figure 7.**
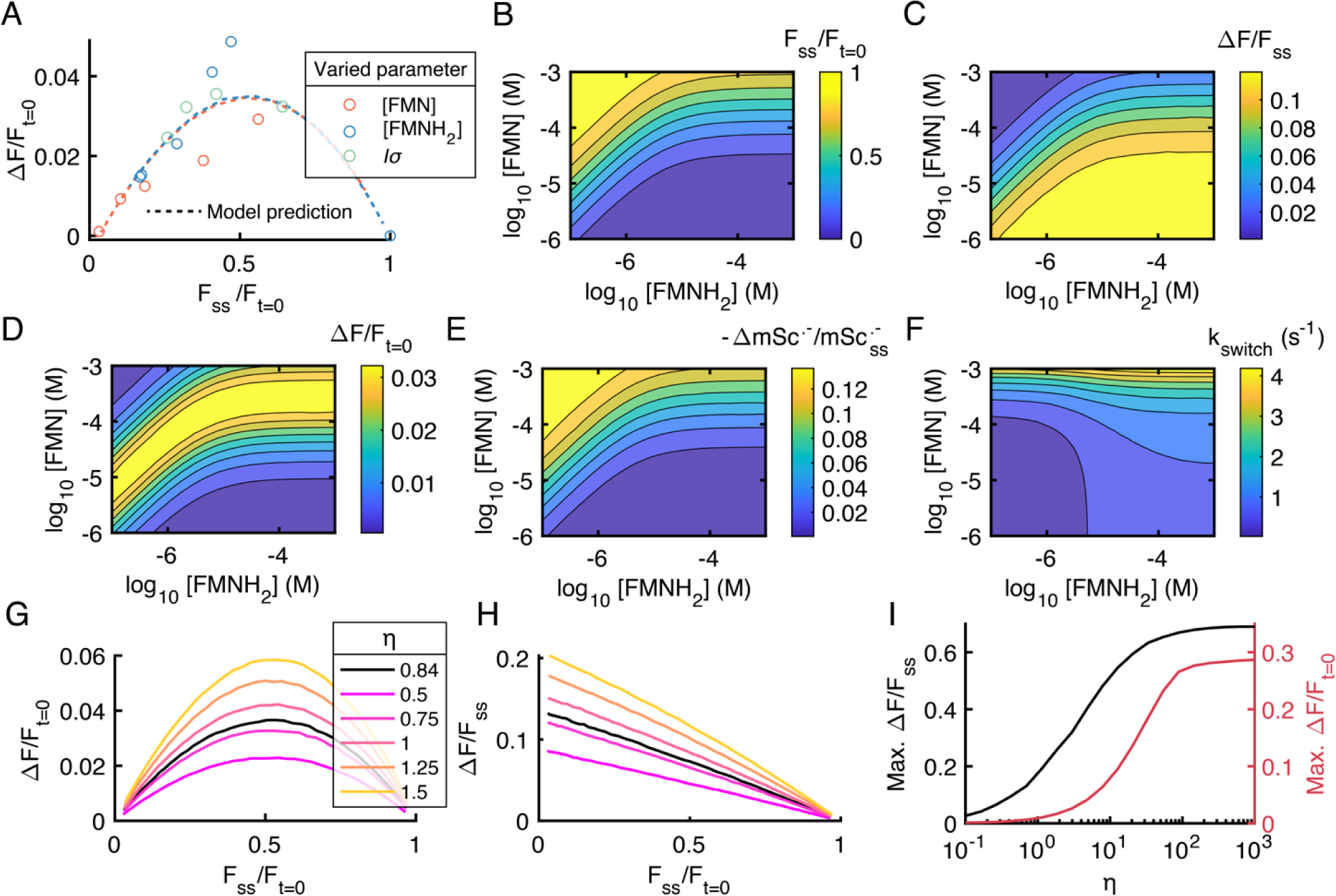
Model predictions for the MFE. A) Universal curve predicts relation between absolute MFE, ΔF/F_t=0_, and residual steady-state fluorescence, x = F_ss_/F_t=0_. Variation in [FMN], [FMNH_2_] and light intensity all led to variations in MFE along this curve. B - F) Predicted effects of [FMNH_2_] and [FMN] on B) F_ss_/F_t=0_, C) ΔF/F_ss_, D) ΔF/F_t=0_, E) 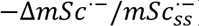, and F) *k*_switch_ for the MFE upstroke. Illumination intensity was assumed to be *Iσ* = 1652 s^-1^, corresponding to 1.7 mW/cm^2^ at 561 nm. G – H) Universal curves for absolute and relative MFE, at several values of the branching ratio *ƞ* = *k*_*rec*_ /*k*_*sep*_. Lines represent represent exact numerical results. [FMN] = 1 – 1000 µM, [FMNH_2_] = 10 µM, *Iσ* = 1652 s^-1^. Analytical approximations in Eqs. 1 and 2. I) Dependence of maximal MFE amplitude on *η*. [FMN] = 0.1 – 100 µM, [FMNH_2_] = 100 µM, *Iσ* = 1652 s^-1^.

The same analysis predicted that the relative MFE followed Δ*F*/*F*_*ss*_ ∝ (1 − *x*), i.e. the relative MFE is largest when the fluorescence is most strongly quenched (*x* ≈ 0). In our experiments in *E. coli*, the steady-state residual fluorescence ranged from *x* = 2.6% at *I* = 0.3 W/cm^2^ to *x* = 0.6% at *I* = 21 W/cm^2^, explaining why we did not detect the intensity-dependent change in Δ*F*/*F*_*ss*_. This analysis presumes that there is no fluorescent background. Presence of a small fluorescent background shifts the maximal value of Δ*F*/*F*_*ss*_ to higher *x*, while having little effect on the maximal value of Δ*F*/*F*_*t*=0_.

The different conditions for maximizing absolute vs. relative MFE highlight a general fact: different chemical quantities may have their magnetic field sensitivity maximized under different conditions. We used the model to predict the time-courses of both observed and unobserved quantities (Fig. S9). Figs. 7B – F show the dependence on [FMN] and [FMNH_2_] of *F*_*ss*_/*F*_*t*=0_, Δ*F*/*F*_*ss*_, Δ*F*/*F*_*t*=0_, the fractional change in mScarlet3^•-^ 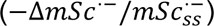, and the magnetic field-induced switching rate, *k*_switch_. Figs. S10 – S11 show the dependence of the same variables on the excitation rate, *Is*, and either [FMN] or [FMNH_2_], and Fig. S12 shows the dependence on [FMN] and *k*_rec_/*k*_sep_.

Finally, we explored the dependence of the universal curves on the parameter *ƞ* ≡ *k*_rec_/*k*_sep_. In the limit of small Δα, the MFE amplitude as function of the steady-state quenched fraction, *x*, is:

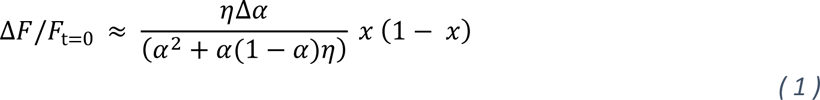

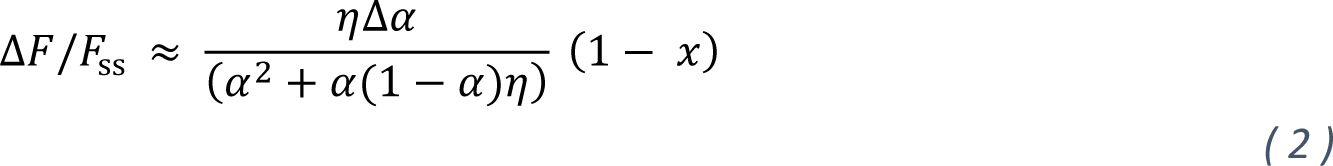

The derivation is in the Supplementary Information. We calculated these quantities from the full numerical model as a function of *x* for different values of *ƞ* (Fig. 7G, H). The numerical model agreed well with these analytical approximations over reasonable parameter ranges.

The maximal MFE across all flavin concentrations and light intensities is determined by the prefactor 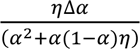 (Fig. 7I). This prefactor is an increasing function of *η*, implying that the MFE is maximized when recombination is faster than separation. In the large-*η* limit, the prefactor becomes 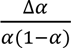, which favors *α* close to 0 or 1, and Δα as large as possible. In summary, to increase the MFE, one should 1) determine which quantity one wishes to maximize, 2) engineer either the quantum mechanical property *α*(B) or the kinetic property *ƞ*, and 3) tune the light intensity, [FMN], and [FMNH_2_] to achieve the requisite amount of quenching.

One may wish to use the MFE to detect time-dependent magnetic fields. The switching rate, *k*_switch_, is the sum of the rates into and out of the mSc^•-^ state. Under continuous low-intensity illumination, *k*_switch_ depends linearly on *Iσ* and [FMN]. Increasing [FMNH_2_] increases *k*_switch_ up until [FMNH_2_] ≈ 5.5 μM. At high intensity (*I* > 10^6^ W/cm^2^), the MFE switching follows a multiexponential, with components corresponding to *k*_ISC_ and *k*_red_[FMNH_2_]. The highest switching rate in our simulations was 1.16 × 10^5^ s^-1^ (Fig. S13), though it is possible our model could fail at such high light intensities, where other photoreactions might come into play.

An alternate approach to measuring fast (and periodic) magnetic field dynamics is to pulse the illumination in synchrony with the oscillations in magnetic field, and to measure the rate of accumulation of mSc^•-^, i.e. as a light-gated integrator. In this case, one should set [FMN] = 0 so that the mSc^•-^ is long-lived. The time-resolution is set simply by the rate of reaction between ^3^mSc and FMNH_2_ to form the SCRP, presuming that the subsequent dynamics are fast. At close-to-saturating [FMNH_2_] = 1.4 mM^23,24^, this rate is predicted to be 7 × 10^4^ s^-1^, implying the ability to use this “lock-in” scheme to detect magnetic fields at up to tens of kHz. The maximum concentration of FMNH_2_ might be increased via addition of hydrotropic agents such as nicotinamide^25,26^, though it is not known how well our model will work at concentrations well outside the range used to determine its parameters.

## Discussion

We determined the mechanism by which small magnetic fields can modulate the fluorescence of mScarlet3. The reaction requires the simultaneous presence of fully reduced FMNH_2_ and oxidized FMN, and involves a photocycle comprising reduction of triplet-state mScarlet3 by FMNH_2_, and re-oxidation of mScarlet3^•-^ by FMN. A numerical model quantitatively reproduced the time-dependent fluorescence trajectories under varied conditions of FMN, FMNH_2_, illumination intensity, and magnetic field, both *in vitro* and in live *E. coli*. We infer that live *E. coli* cytoplasm contains sufficient FMN and FMNH_2_ (or perhaps other oxidant and reductant), so the MFE does not require blue light. Previous work has shown that other redox cofactors such as cinnavalininate and WST-8 can also enable the MFE^1^.

Some molecular details in our model remain uncertain. The molecular structure of the putative mScarlet3^•-^ is not known. Many red fluorescent proteins — KillerRed, mRFP, DsRed, and mKate2 — accept electrons upon photo-excitation^27–29^, suggesting the presence of a common mechanism. Electron spin resonance (ESR) studies may provide insights into the nature of the radical anion. The detailed mechanism underlying electron-transfer between FMNH_2_ and ^3^mSc is also unknown. Fluorescent proteins undergo multiple light-driven redox reactions, some involving direct electron-transfer between the chromophore and an external redox species, and some involving hopping mediated by other amino acids in the protein.^29^ Further mutagenesis, transient absorption, and computational studies will be needed to elucidate the details of this reaction step.

In organic donor-acceptor complexes, the MFE can be enhanced by tethering the groups together,^5,30^ or by confinement in micelles^10,31^. In the context of our model, these effects can be understood as increasing the parameter *η*. Thus, tethering the flavin cofactor or nano-confinement (in micelles, vesicles, or pores in a gel) may substantially enhance the MFE amplitude. Tethering a fluorescent protein to a flavin-binding domain may enable development of magneto-switchable proteins in mammalian cells, a necessary step for opaque-tissue applications of magnetogenetics.

To increase the magnetic field sensitivity, one should decrease the strength of the hyperfine coupling at the locations of the spins. It may be possible to identify redox cofactors that have weaker hyperfine couplings than FMNH^•-^; whether directed evolution can modulate the hyperfine environment for the electron spin on mSc^•-^ remains an open question.

A detailed mechanistic understanding of spin states in fluorescent proteins may lead to new microscopy and sensing techniques.^32^ For example, a recent study demonstrate fluorescence-detected electron-spin resonance via optical pumping of the triplet state in yellow fluorescent protein (YFP)^33^. Another recent study showed fluorescence-detected electron-spin resonance in a modified LOV domain, magLOV^34^. In organic solutions showing magnetic-sensitive fluorescence, imaging through a scattering medium was achieved by sweeping a magnetic null through the sample^35^. Development of protein-based spintronics requires a detailed understanding of both the quantum spin dynamics, and also the kinetics of the photocycles in which these spin dynamics are embedded.

Photo-redox reactions are common in proteins: cryptochromes and photolyases contain flavin cofactors^36^, and protein backbones can donate electrons to photo-excited FMN^7,36^. Among fluorescent proteins, GFP can donate an electron from the excited singlet state, undergoing oxidative redding.^37,38^ Under anaerobic conditions, GFP may also accept an electron^38–40^. We suspect that many proteins with these properties may be magnetic field-sensitive or could be engineered to show magnetosensitivity under suitable conditions.

It remains controversial whether radical pair-based magnetic field effects have any biological function^41,42^. The half-maximal field of 5.5 mT in our experiments is approximately 100-fold larger than Earth’s magnetic field, so our experiments do not directly address this controversy. Regardless of whether magnetic field effects serve any physiological function, the existence of strong MFEs in proteins under physiological conditions suggests that such effects may be more widespread than currently appreciated. An exciting future opportunity is to use the power of protein design and directed evolution to tune the MFE amplitude, magnetic field sensitivity, and switching kinetics; and to couple the MFE to downstream biochemical cascades.

## Methods

### Bacterial culture and sample preparation

6xHis-tagged pDx_mScarlet3 (Addgene #189754) was expressed in BL21 (DE3) *E. coli* (NEB C2527H) and plated on lysogeny broth (LB) agar plates with 100 µg/mL Kanamycin (Teknova L1025). The plasmid sequence was verified through whole-plasmid sequencing (Plasmidsaurus). Bacteria were grown at 37 °C overnight. The next day, a single colony was inoculated in 5 mL of LB supplemented with 50 µg/mL Kanamycin and grown in a shaking incubator at 37 °C and 250 r.p.m. overnight.

250 µL from the liquid culture was used to inoculate a 25 mL culture. After 1 hour in a shaking incubator at 37 °C, L-Rhamnose (Promega L5701) was added to a final concentration of 0.2% (w/v) to induce mScarlet3 expression. Bacteria were grown for 4 additional hours to an OD700 of ∼0.4. The culture was stored overnight at 4 °C. The liquid culture was imaged the next day.

Media was aspirated and 2 µL of the wet pellet was mounted between a glass slide and a #1.5 coverslip and sealed with clear nail polish.

For the initial screening for MFEs (Fig. S1), samples were grown uninduced on LB agar plates with the appropriate selection antibiotic. The additional fluorescent proteins tested were mScarlet-I3 (Addgene #189757), mRuby3 (Addgene #104005), mSandy2 (Addgene #177760), and mKate2 (Addgene #104030).

### Protein purification

His-tagged mScarlet3 was purified using Ni-NTA affinity columns and the kit protocol and buffers (Qiagen 30600). A 10 mL overnight culture was used to inoculate a 250 mL culture. After 1 hour in a shaking incubator at 37 °C, L-Rhamnose (Promega L5701) was added to a final concentration of 0.2% (w/v) to induce mScarlet3 expression. Bacteria were grown for 4 additional hours, and then stored overnight at 4 °C. Cells were harvested by centrifugation at 3400 g for 20 minutes and frozen at -20 °C overnight. The pellet was thawed for 15 minutes on ice and resuspended in 10 mL native Lysis Buffer with one tablet of cOmplete™ Protease Inhibitor Cocktail (Roche 11836170001). The cell lysate was centrifuged at 14,000 g for 30 minutes at 4 °C, and the supernatant was applied to the Fast Start Column. The column was washed twice with 4 mL of Native Wash Buffer. The protein was eluted in two 1 mL fractions of Native Elution Buffer. Imidazole was removed via by spin column filtration. Samples were measured in 1x PBS buffer. Concentrations were quantified using a Nanodrop, using *ε*_570_ = 104,000 M^-1^ cm^-1^. For immobilized mScarlet3 experiments, agar Ni-NTA beads from the purification resin were washed with PBS and imaged.

### Fluorescence measurements

Samples were imaged on a home-built inverted epifluorescence microscope with 488 nm, 532 nm, and 561 nm laser lines. The beams were combined using dichroic mirrors and sent through an acousto-optic tunable filter (AOTF; Gooch and Housego TF525-250-6-3-GH18A) for temporal modulation of intensity of each wavelength. The beams were expanded and focused to the back-focal plane of an objective (Olympus 20X/1.00 W XLUMPLANFL, or Olympus 40x/0.85 UPlanApo) installed in an Olympus IX71 microscope. None of the elements of the sample stage was magnetic.

Excitation light was separated from fluorescence emission with a 562 nm edge dichroic mirror (Semrock Di02-FF562) and a 575 nm longpass emission filter (Chroma ET575lp). In experiments where FMN fluorescence was monitored, a quad-edge dichroic (Semrock Di01-405/488/561/635) was used with a dual band emission filter (Chroma ZET488/561m). Emission light was sent to an imaging spectrometer (Horiba iHR320) equipped with an EMCCD camera (Andor iXon EM+). For imaging experiments, the zeroth order was taken from the spectrometer and the slit was set to the maximum width of 60 mm, allowing the full image to pass through to the camera. For measurements of fluorescence emission spectra, we calibrated the linear pixel-to-wavelength correspondence by sending the laser lines directly through using a 50/50 beamsplitter. We used a 1–2 mm entrance slit width for measurements on the microscope. Data were five-point moving average filtered.

Data in Fig. 3 were taken using the same excitation lasers. The zeroth order from the AOTF was expanded to illuminate a quartz cuvette (Starna Cells 23-G-5). A mirror on one side of the cuvette were used to achieve two 10 mm passes of the excitation light through the cuvette. The emission was collected perpendicular to the excitation using a fiber and collimated into the same imaging spectrometer with a 2 mm entrance slit width. Excitation lasers were temporally modulated using physical shutters.

Magnetic fields were modulated by using a permanent magnet (K&J Magnetics B666) mounted on a servo motor. The distance from the sample was adjusted to control the field strength. Square wave magnetic fields were applied with a ∼200 ms rise time, with field strength and timing measured by a Hall effect sensor (Texas Instruments DRV5055A4QLPG).

The AOTF, shutters, servo, and the camera were synchronized via a National Instruments Data Acquisition System (DAQ) and custom software (https://www.luminosmicroscopy.com/).

All measurements were performed at 22 °C (295 K).

### Absorption measurements

Absorption spectra were taken with a path length of 5 mm (Starna Cells 23-G-5) using a UV-VIS-NIR lamp (Mikropack DH-2000). The same fiber was used to collect transmitted light and fluorescence emission. The transmission light filled the volume of the sample. Physical shutters for the transmitted light and for the excitation light were modulated separately. The transmission light used for absorption measurements was dim, to avoid exciting fluorescence or driving photoswitching.

## Acknowledgments

We thank Madeleine Howell, Daniel G. Itkis, Xiang Wu, F. Phil Brooks III, Peter Maurer, Uri Zvi, Ben Soloway, and Jacob Feder for helpful discussions and technical assistance. This work was supported by the Gordon and Betty Moore Foundation and NSF Quantum Sensing for Biophysics and Bioengineering (QuBBE) Quantum Leap Challenge Institute (QLCI) grant OMA-2121044. K. M. X. was supported by a Hertz Foundation Fellowship and an NSF Graduate Research Fellowship.

## Supplementary Figures

**Figure S1.**
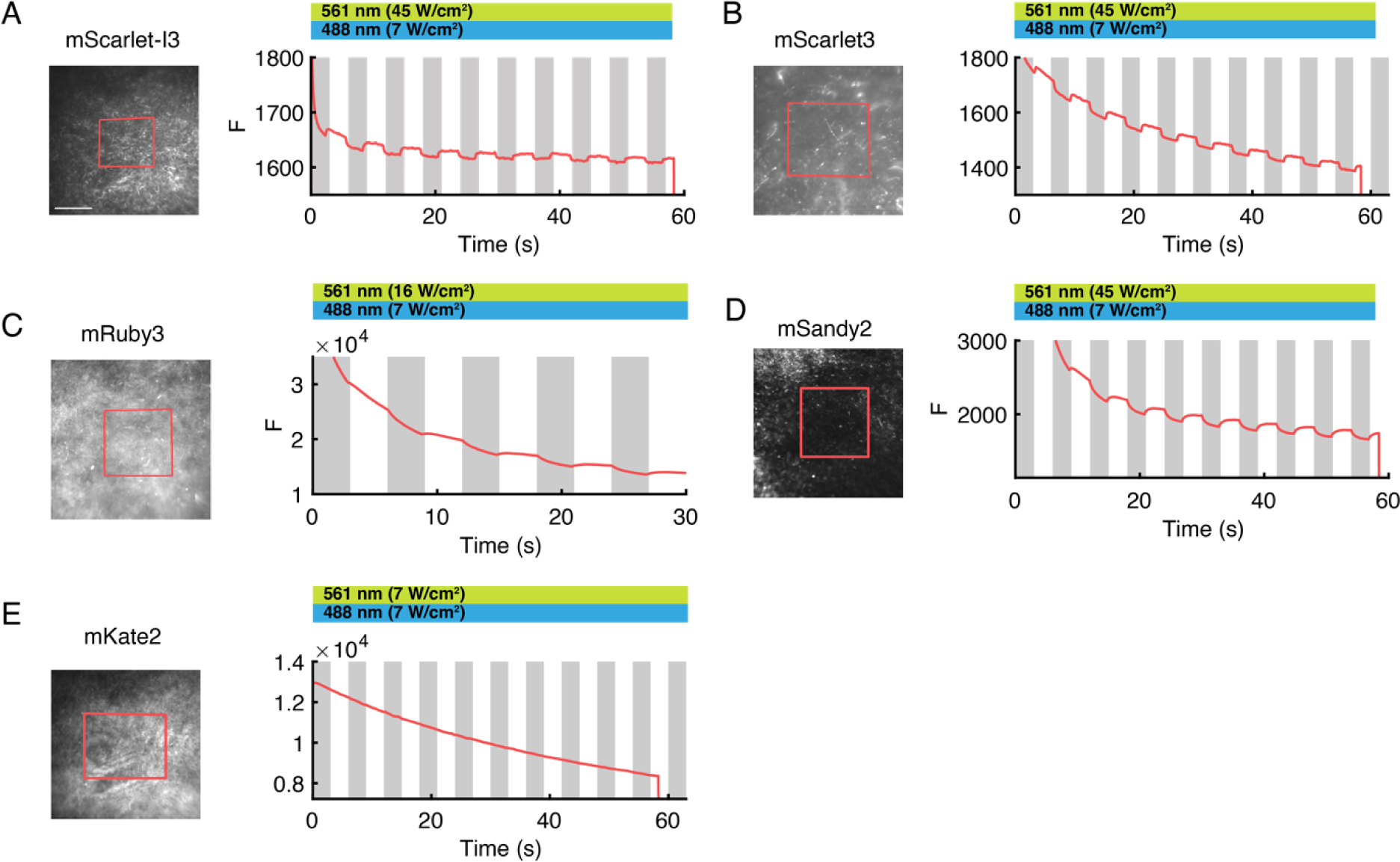
Magnetic field effects in red fluorescent proteins. *E. coli* expressing red-emitting fluorescent proteins were grown on LB (lysogeny broth) agar plates and then imaged in an epifluorescence microscope. Panels show responses to a 10 mT magnetic field (grey bars). A 595 nm longpass emission filter was used. A) mScarlet-I3,^11^ B) mScarlet3,^11^ C) mRuby3,^43^ D) mSandy2,^44^ E) mKate2.^45^

**Figure S2.**
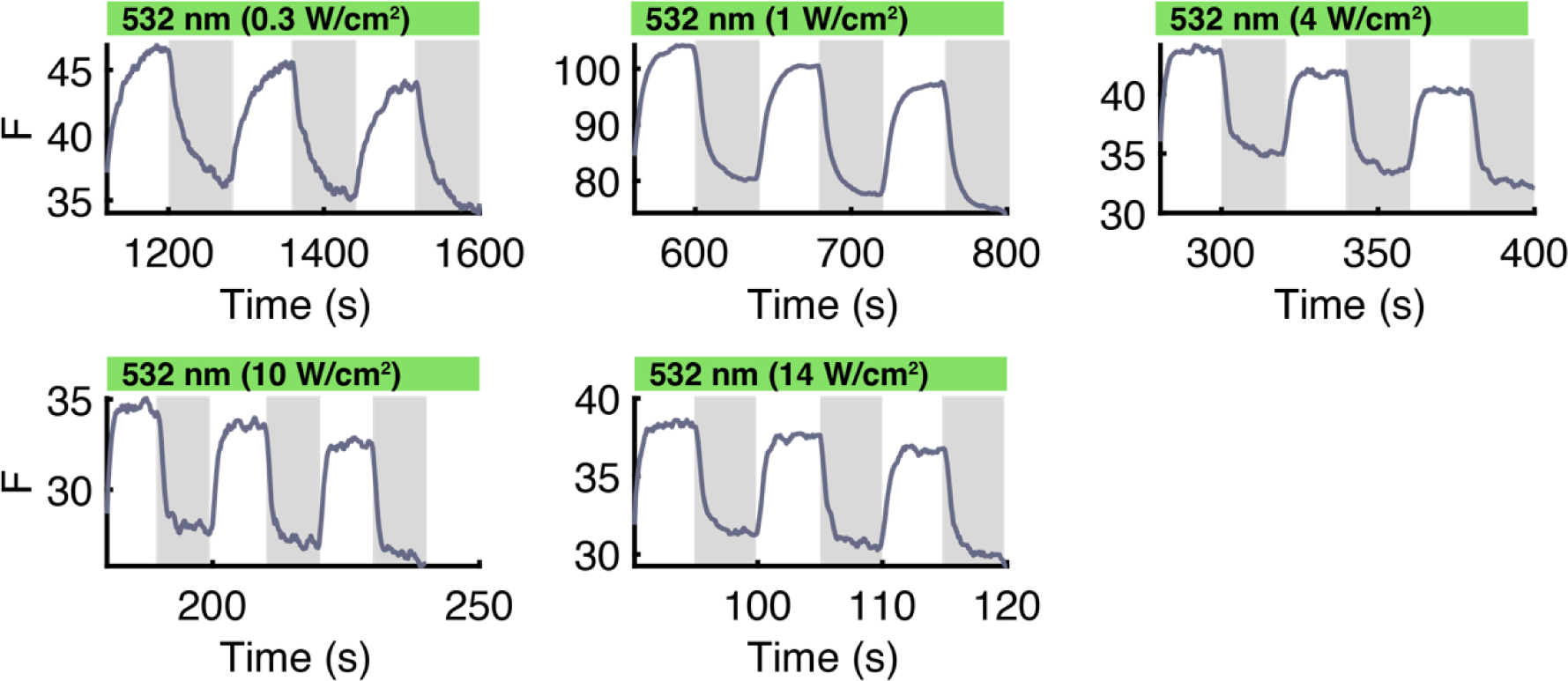
MFE kinetics depend on light intensity. MFE switching kinetics in *E. coli* at different illumination intensities. Plots show fluorescence after initial fluorescence has quenched to a steady-state.

**Figure S3.**
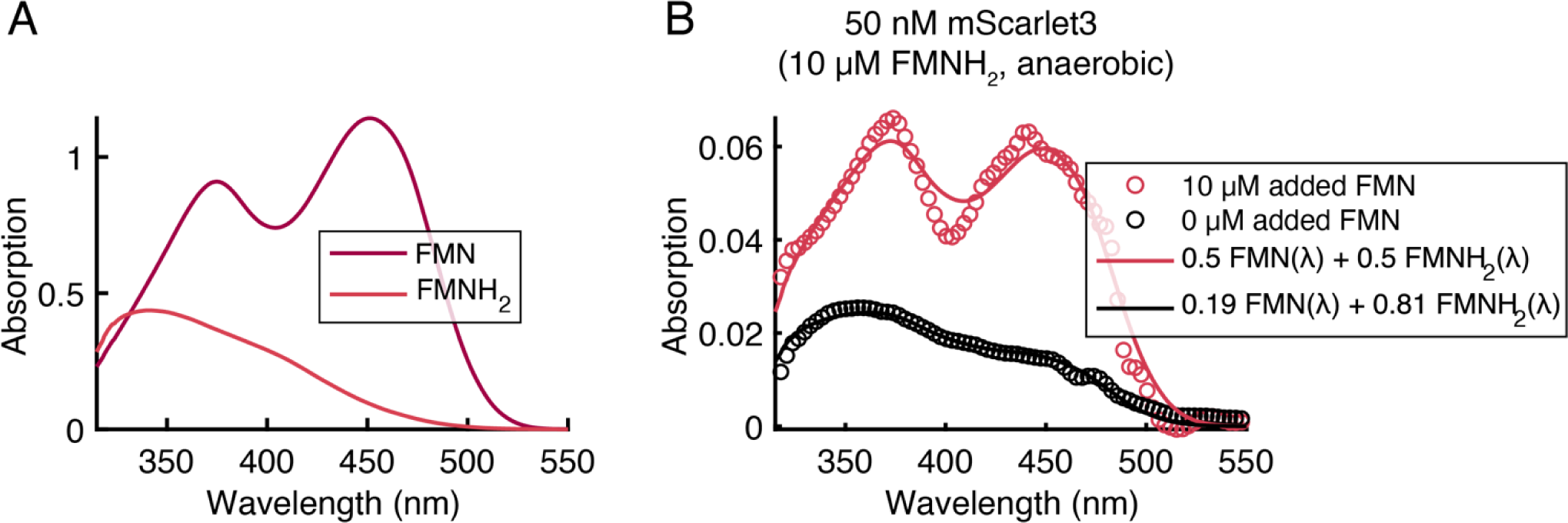
Flavin absorption spectra. A) Absorption spectra of pure FMN and FMNH_2_. The spectrum for FMNH_2_ was measured immediately after photoreduction in the presence of 50 mM EDTA under anaerobic conditions. B) Absorption spectra for the samples in Fig. 4G, comprising 10 µM of photoreduced FMN (nominally FMNH_2_) and either 10 µM or 0 µM additional FMN. The black curve shows that the nominal 10 µM FMNH_2_ actually contained 2 µM FMN, due to incomplete photoreduction.

**Figure S4.**
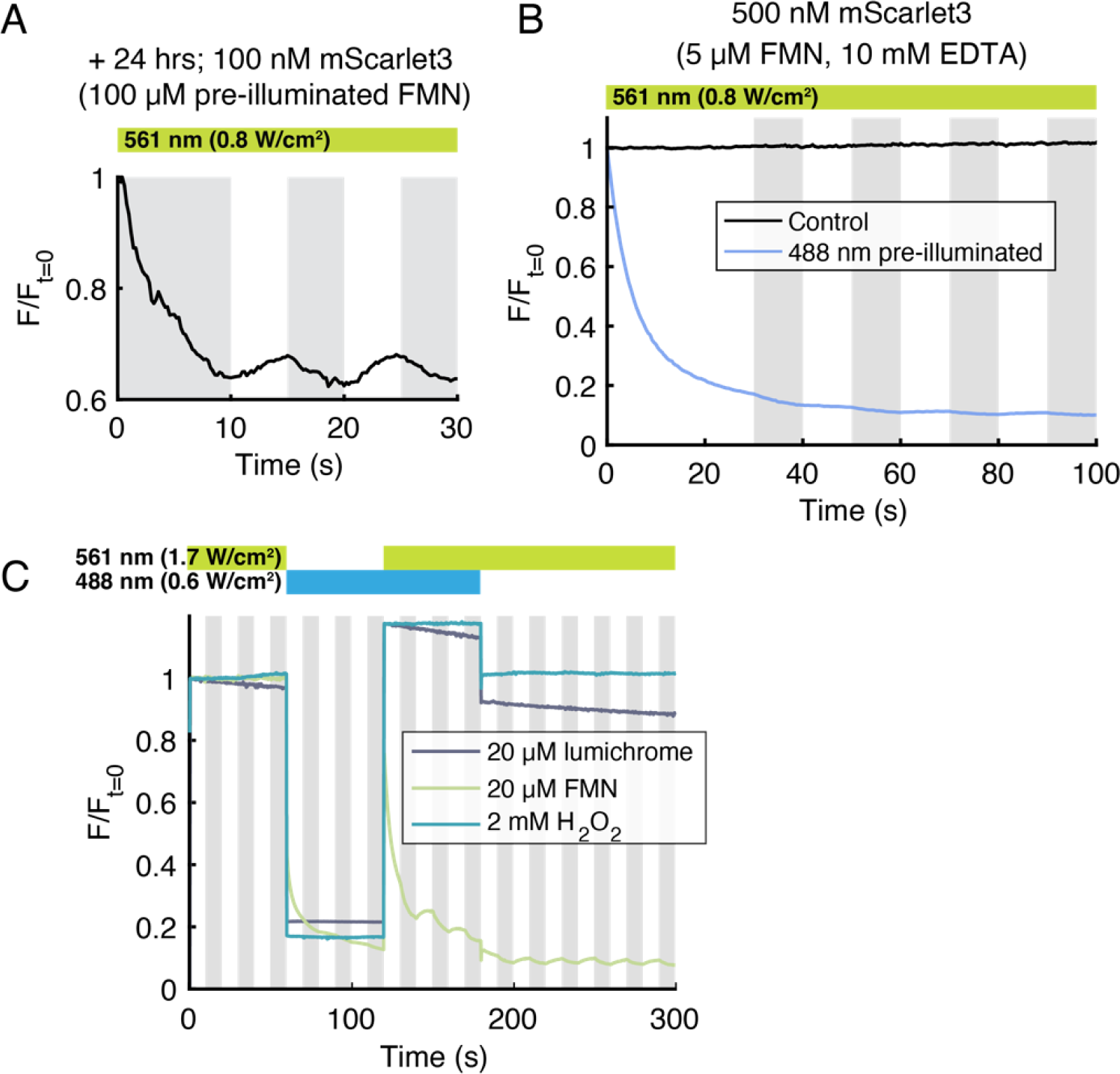
Control experiments confirming FMNH_2_ is the redox-active FMN photoproduct. A) FMNH_2_ was photochemically produced by illuminating 100 mM FMN with 488 nm light, under anaerobic conditions. The FMNH_2_ was then combined with mScarlet3 and then stored at room temperature in the dark under anaerobic conditions for 24 h. Under yellow light alone, the sample showed a clear MFE, indicating the stability of the FMN photoproduct under these conditions. B) mScarlet3, FMN, and 10 mM EDTA were combined under anaerobic conditions. The sample showed no quenching or MFE upon 561 nm illumination, indicating that EDTA does not reduce FMN under these conditions. The sample was then pre-illuminated with 488 nm light. Upon onset of yellow illumination, the mScarlet fluorescence quenched and showed an MFE, as expected. C) mScarlet3 did not show an MFE when combined with other FMN photoproducts including lumichrome (mScarlet3 at 500 nM) or H_2_O_2_ (mScarlet3 at 5 µM), under either 488 nm, 561 nm, or combined 488 + 561 nm illumination. Trace with 500 nM mScarlet3 and 20 µM FMN shown for comparison. All samples were measured under anaerobic conditions.

**Figure S5.**
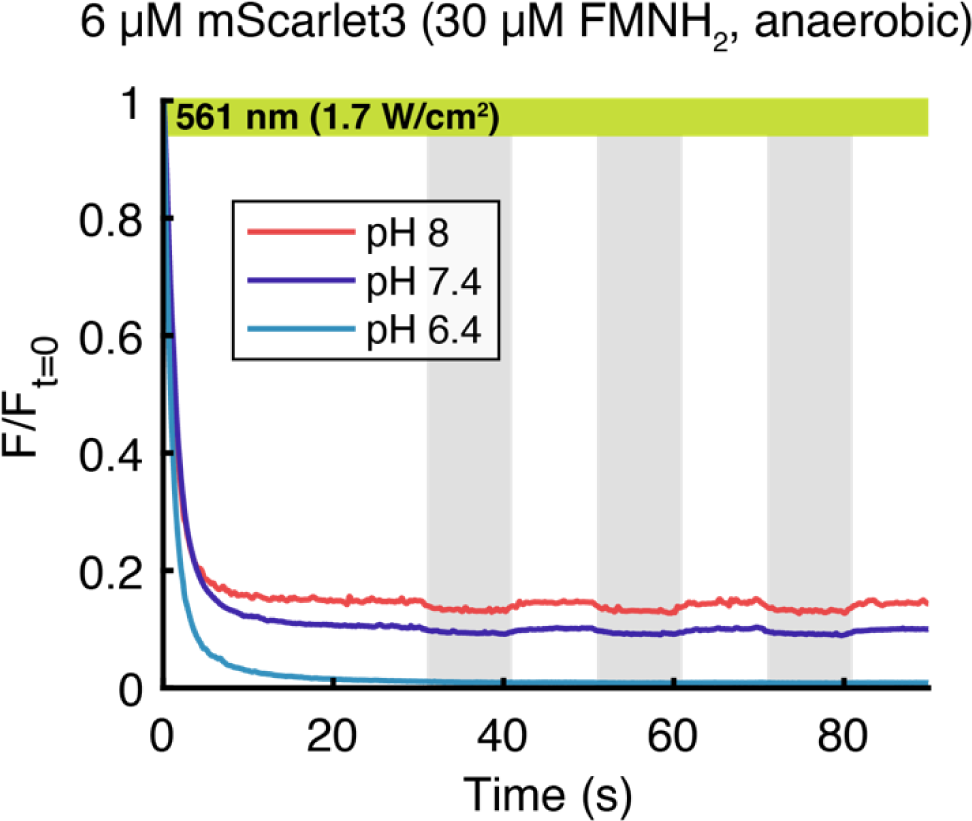
pH dependence of mScarlet3 fluorescence quenching and recovery.

**Figure S6.**
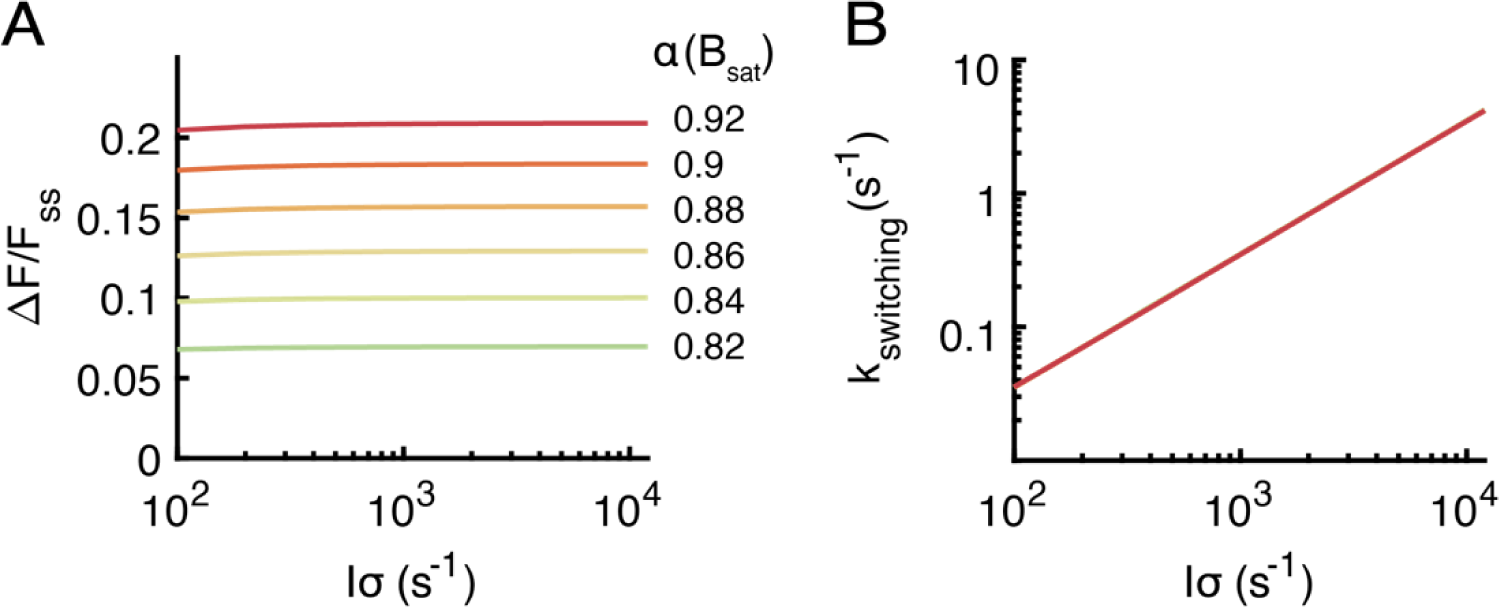
Effect of varying the SCRP triplet fraction at saturating magnetic field. Numerical predictions of A) MFE amplitude (compare to Fig. 1F), and B) MFE switching kinetics (compare to Fig. 1H). For these simulations, we used 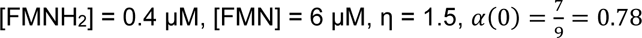.

**Figure S7.**
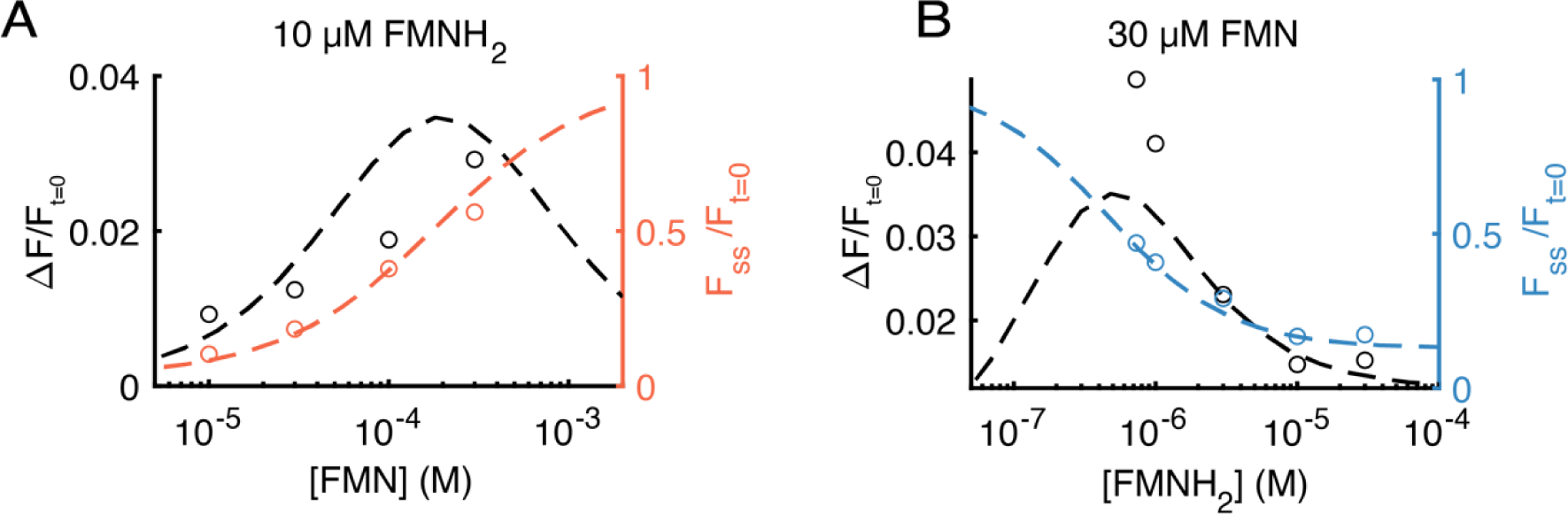
Model predictions of flavin concentration dependence. Numerical model prediction (dashed lines) for the absolute MFE (data: black circles) and the fractional quenching (data: orange and blue circles) as a function of the concentration of A) FMN titrated at 10 µM FMNH_2_, B) FMNH_2_ titrated at 30 µM FMN. Data are from Figs. 4G, H.

**Figure S8.**
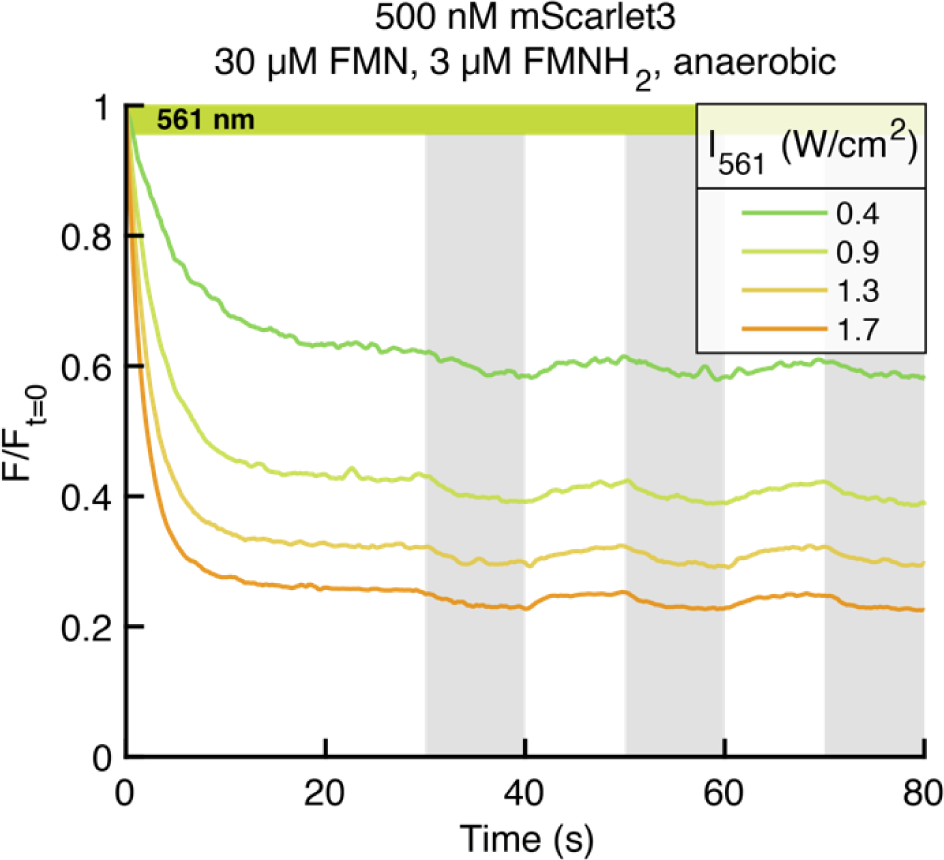
Effect of illumination intensity on MFE in purified mScarlet3. Higher intensity light led to larger initial quenching, faster MFE switching, and larger relative MFE, ΔF/F_ss_. Samples were prepared under the same conditions as in Figs. 4G, H.

**Figure S9.**
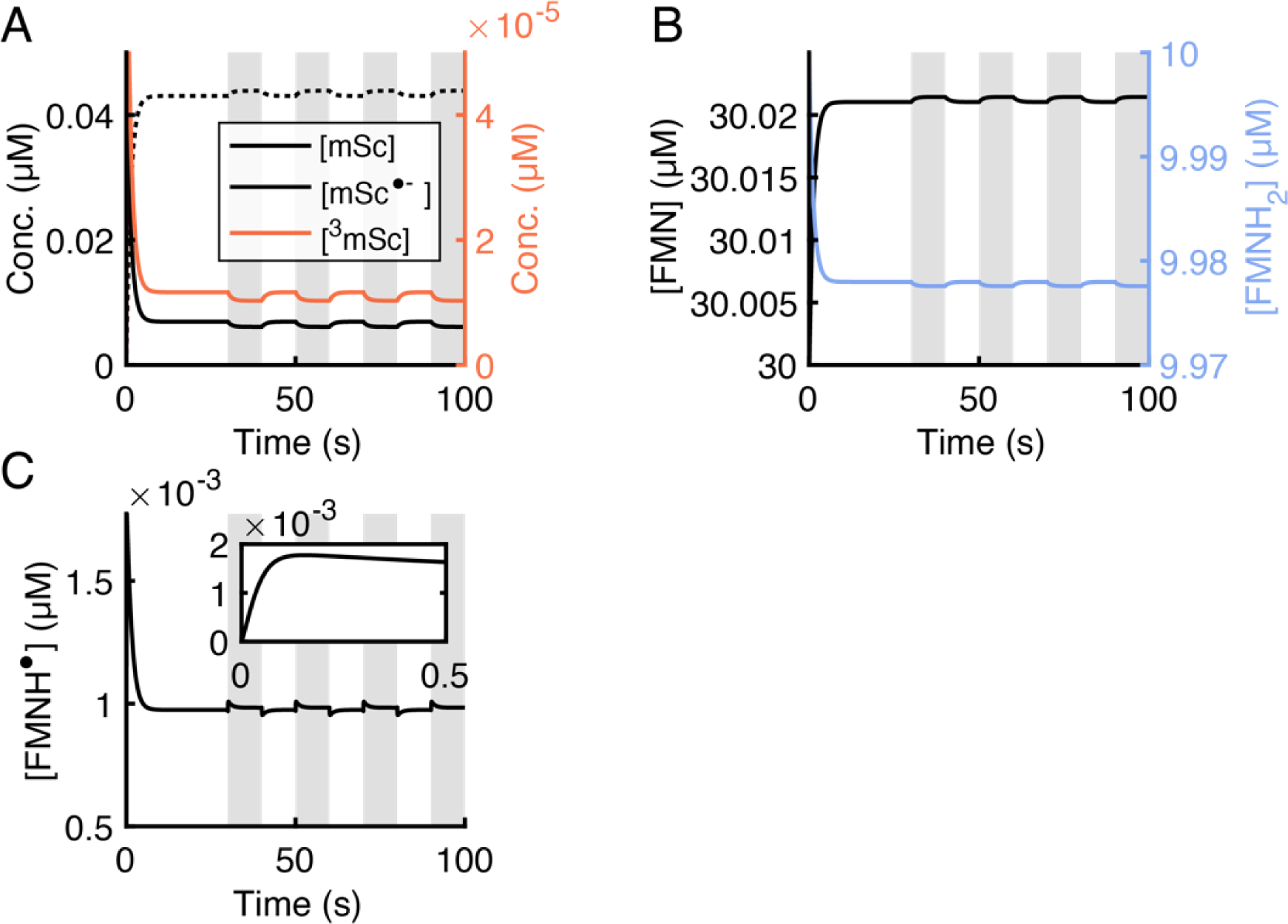
MFE in photocycle intermediates. Simulated kinetic traces of mScarlet3 and flavin species. Initial mScarlet3 at 50 nM, FMN at 30 µM, FMNH_2_ at 10 µM. A) Upon onset of mScarlet3 photo-excitation (*I*σ = 1652 s^-1^, 1.7 W/cm^2^), ground state mScarlet3 rapidly quenches while mSc^•-^ increases. Most mScarlet3 is in these two states; here 0.1% is in the triplet state, ^3^mSc. B) FMN and FMNH_2_ concentrations reach a steady-state on the same timescale as the quenching, and show a small MFE since they are in excess of mScarlet3. C) FMNH^•^ concentration shows a small two-component response to the magnetic field. Inset: fast initial dynamics of FMNH^•^ upon production and subsequent separation of the SCRP.

**Figure S10.**
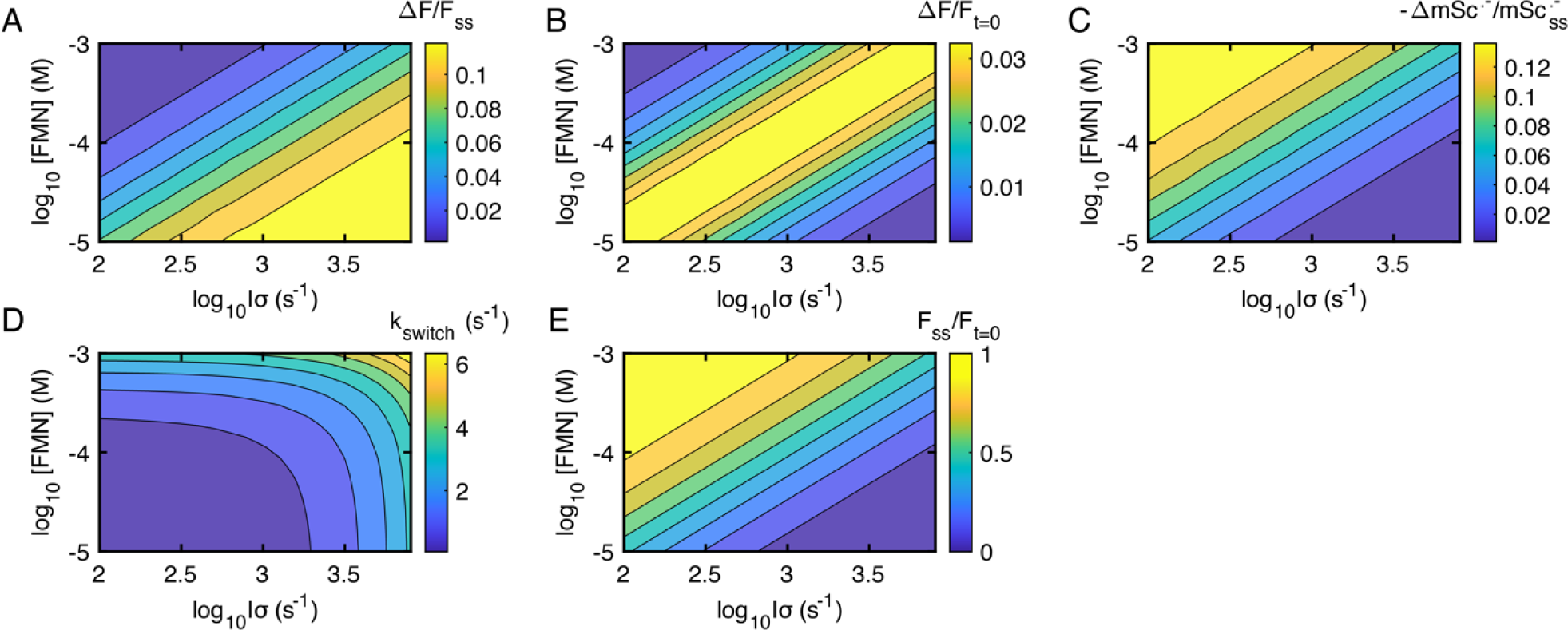
Dependence of MFE photophysics on illumination intensity and [FMN]. Contour plots of A) ΔF/F_ss_, B) ΔF/F_t=0_, C) -ΔmSc^•-^/mSc^•-^_0_, D) k_switch_ for the MFE upstroke, and E) F_ss_/F_t=0_ as a function of [FMN] and light intensity. Here [FMNH_2_] = 10 µM.

**Figure S11.**
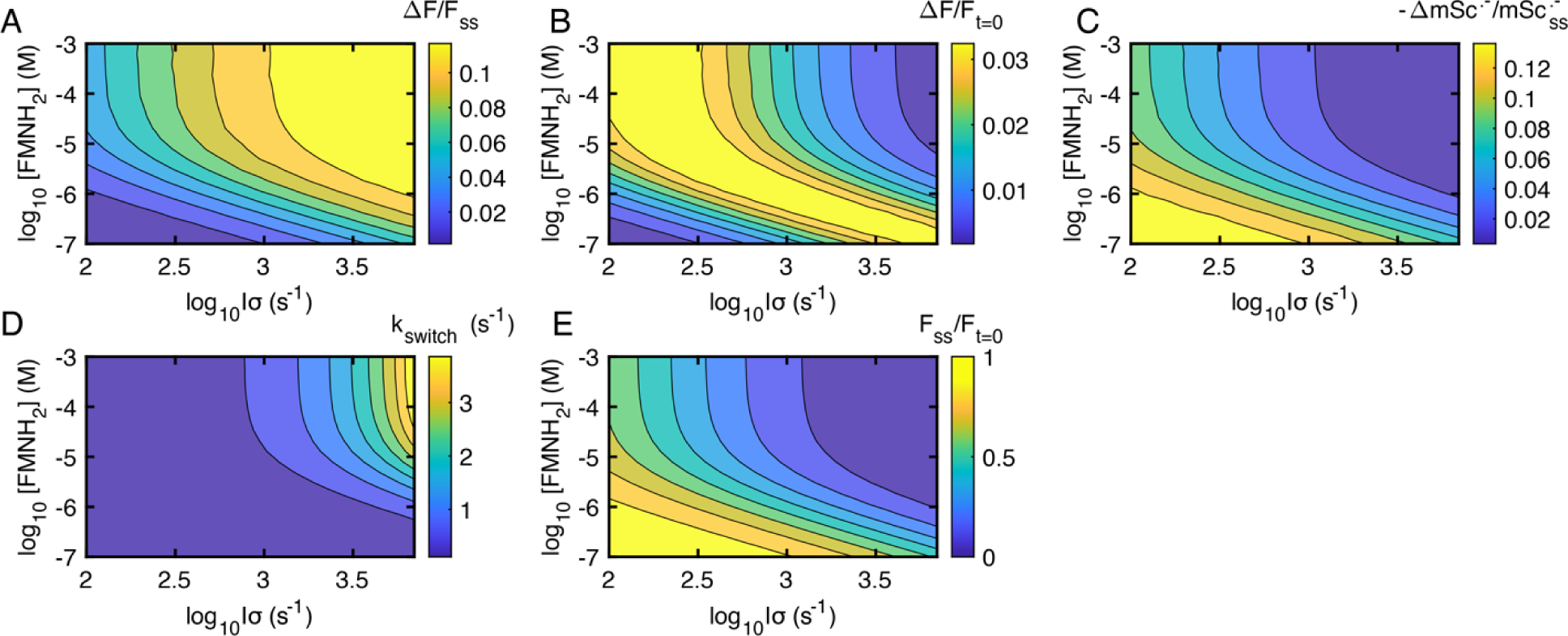
Dependence of MFE photophysics on illumination intensity and [FMNH_2_]. Contour plots of A) ΔF/F_ss_, B) ΔF/F_t=0_, C) -ΔmSc^•-^/mSc^•-^_0_, D) k_switch_ for the MFE upstroke, and E) F_ss_/F_t=0_ as a function of [FMNH_2_] and light intensity. Here [FMN] = 30 µM.

**Figure S12.**
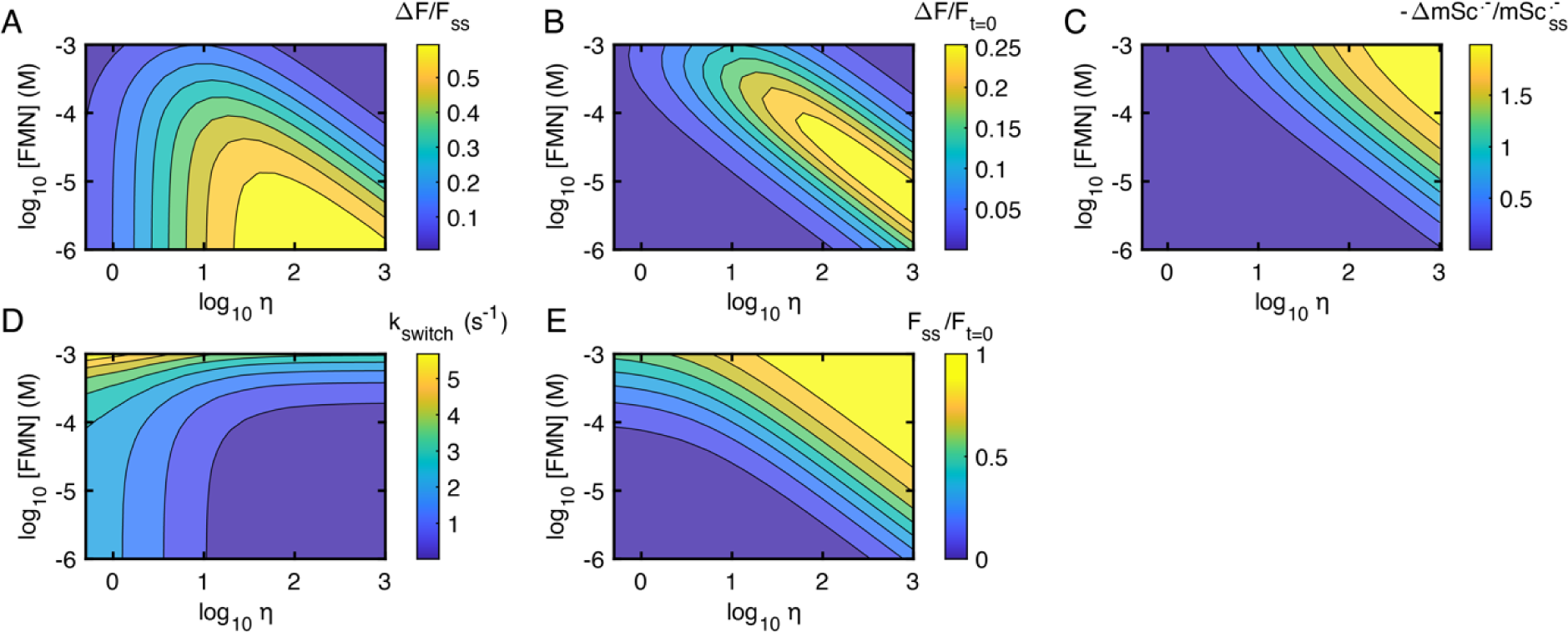
Dependence of MFE photophysics on *η* and [FMN]. Contour plots of A) ΔF/F_ss_, B) ΔF/F_t=0_, C) -ΔmSc^•-^/mSc^•-^_0_, D) k_switch_ for the MFE upstroke, and E) F_ss_/F_t=0_ as a function of [FMN] and η = k_rec_/k_sep_. Here [FMNH_2_] = 10 µM, *Iσ* = 1652 s^-1^.

**Figure S13.**
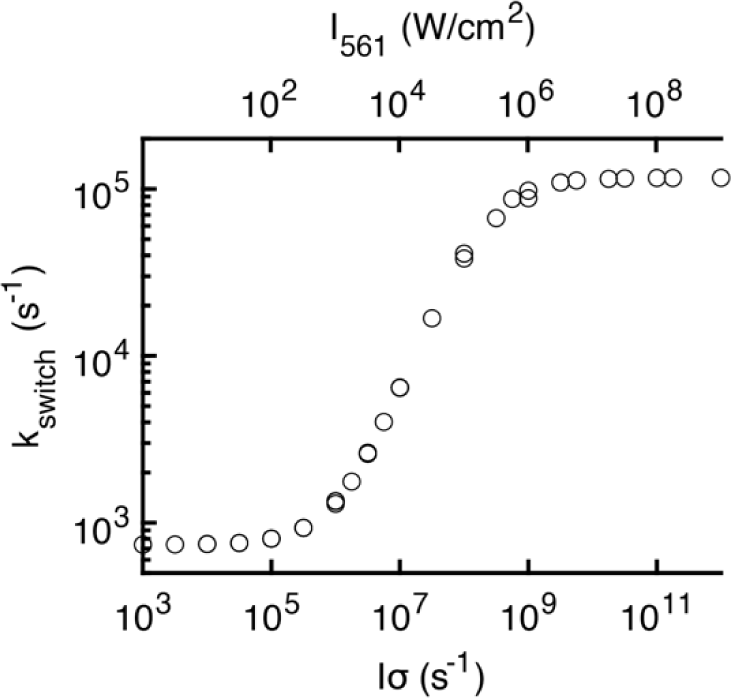
Dependence of predicted MFE switching rate k_switch_ on high light intensities. The MFE switching rate (fit to an exponential + constant) increases with light intensity until 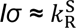. Here, [FMN] = 200 mM and [FMNH_2_] = 1.4 mM, which are the predicted maximal solubilities in water^23,24^.

