## Supplementary Information for "Mechanism of giant magnetic field effect in fluorescence of mScarlet3, a red fluorescent protein"

### Kinetic modeling of the MFE in mScarlet3

#### 1 Photocycle of mScarlet3/FMN/$FMNH_2$

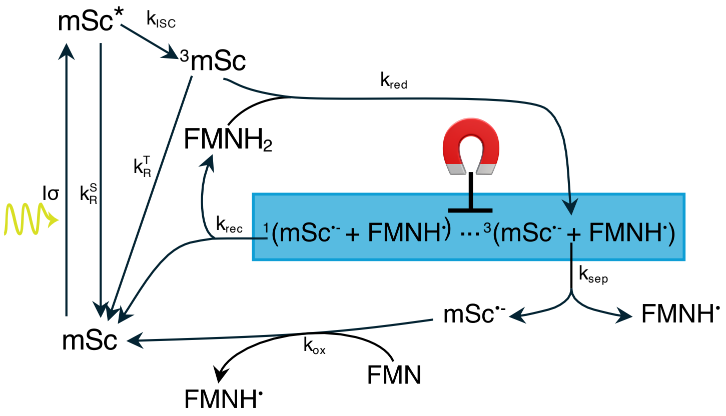


Photocycle of mScarlet3 in the presence of FMN and $FMNH_2$.

The following chemical reactions describe the model:

$$\text{mSc} + h\nu \underset{k_R^S}{\overset{I\sigma}{\rightleftarrows}} \text{mSc}^* \tag{1}$$

$$\text{mSc}^* \xrightarrow{k_{ISC}} {}^3\text{mSc} \tag{2}$$

$${}^3\text{mSc} \xrightarrow{k_R^T} \text{mSc} \tag{3}$$

$${}^3\text{mSc} + \text{FMNH}_2 \xrightarrow{k_{red}} \text{SCRP} \tag{4}$$

$$\alpha\text{SCRP} \xrightarrow{k_{sep}} \text{mSc}^{\bullet-} + \text{FMNH}^{\bullet} \tag{5}$$

$$(1\text{-}\alpha)\text{SCRP} \xrightarrow{k_{rec}} \text{mSc} + \text{FMNH}_2 \tag{6}$$

$$\text{mSc}^{\bullet-} + \text{FMN} \xrightarrow{k_{ox}} \text{mSc} + \text{FMNH}^{\bullet} \tag{7}$$

$$\text{FMNH}^{\bullet} + \text{FMNH}^{\bullet} \xrightarrow{k_{dis}} \text{FMN} + \text{FMNH}_2 \tag{8}$$

Since we are interested in dynamics on timescales long compared to the [mSc*] lifetime (4 ns), we make the simplifying approximation $\frac{d[\text{mSc}^*]}{dt} \approx 0 \approx I\sigma[\text{mSc}] - (k_R^\text{S} + k_\text{ISC})[\text{mSc}^*]$. Thus, $[\text{mSc}^*] \approx \frac{I\sigma}{k_\text{ISC}+k_R^\text{S}}[\text{mSc}]$, and upon onset of illumination the initial concentration of [mSc] is $[\text{mSc}](t = 0^+) = \left(\frac{k_R^\text{S}+k_\text{ISC}}{k_R^\text{S}+k_\text{ISC}+I\sigma}\right)[\text{mSc}^\text{Tot}]$, where $[\text{mSc}^\text{Tot}]$ is the total concentration of mScarlet3.

Reactions 1 and 2 then simplify to:

$$\text{mSc} + h\nu \underset{k_R^\text{T}}{\overset{k_\text{ISC}^\text{eff}}{\rightleftarrows}} {}^3\text{mSc}, \qquad \{9\}$$

where

$$k_\text{ISC}^\text{eff} = \frac{I\sigma\, k_\text{ISC}}{k_\text{ISC} + k_R^\text{S}}$$

is the effective rate from [mSc] into the triplet state $[^3\text{mSc}]$. The kinetic equations then become:

$$\begin{aligned}
\frac{d[\text{mSc}]}{dt} &= -k_\text{ISC}^\text{eff}[\text{mSc}] + k_R^\text{T}[^3\text{mSc}] + k_\text{rec}(1-\alpha)[\text{SCRP}] + k_\text{ox}[\text{FMN}][\text{mSc}^{\bullet-}] \\
\frac{d[^3\text{mSc}]}{dt} &= +k_\text{ISC}^\text{eff}[\text{mSc}] - \left(k_R^\text{T} + k_\text{red}[\text{FMNH}_2]\right)[^3\text{mSc}] \\
\frac{d[\text{SCRP}]}{dt} &= k_\text{red}[^3\text{mSc}][\text{FMNH}_2] - (\alpha k_\text{sep} + (1-\alpha)k_\text{rec})[\text{SCRP}] \\
\frac{d[\text{mSc}^{\bullet-}]}{dt} &= k_\text{sep}\alpha[\text{SCRP}] - k_\text{ox}[\text{FMN}][\text{mSc}^{\bullet-}] \\
\frac{d[\text{FMN}]}{dt} &= -k_\text{ox}[\text{FMN}][\text{mSc}^{\bullet-}] + k_\text{dis}[\text{FMNH}^\bullet]^2 \\
\frac{d[\text{FMNH}_2]}{dt} &= -k_\text{red}[^3\text{mSc}][\text{FMNH}_2] + k_\text{rec}(1-\alpha)[\text{SCRP}] + k_\text{dis}[\text{FMNH}^\bullet]^2 \\
\frac{d[\text{FMNH}^\bullet]}{dt} &= k_\text{sep}\alpha[\text{SCRP}] - 2k_\text{dis}[\text{FMNH}^\bullet]^2 + k_\text{ox}[\text{FMN}][\text{mSc}^{\bullet-}].
\end{aligned}$$

To simplify further, we next analyze the dynamics in the SCRP state.

### 2 SCRP interconversion approximation

The timescale of interconversion between the singlet and triplet SCRP is approximately $\gamma_e B_{1/2}$, where $\gamma_e =$ 28 MHz/mT is the electron gyromagnetic ratio, and $B_{1/2} = 5.5$ mT is the experimentally measured half-saturating magnetic field – approximately the strength of the nuclear hyperfine fields coupled to each electron in the SCRP. Thus $k_\text{ISC}^\text{SCRP} \approx 10^9$ s$^{-1}$, faster than the rates into and out of the SCRP. Hence in our model we use the magnetic field-dependent branching ratio $0 < \alpha < 1$ to describe the fraction of SCRP in the triplet manifold at quasi-steady state.

We further assume that the SCRP concentration reaches steady-state quickly compared to the dynamics of interest, so we approximate $\frac{d[\text{SCRP}]}{dt} \approx 0$. Reactions 4–6 then become:

$$\frac{d[\text{SCRP}]}{dt} \approx 0 \approx k_\text{red}[\text{FMNH}_2][^3\text{mSc}] - \alpha k_\text{sep}[\text{SCRP}] - (1-\alpha)k_\text{rec}[\text{SCRP}]$$

Therefore, the concentration of the SCRP is proportional to the mScarlet3 triplet:

$$[\text{SCRP}] = \frac{k_{\text{red}}}{\alpha k_{\text{sep}} + (1-\alpha) k_{\text{rec}}} [\text{FMNH}_2][^3\text{mSc}]$$

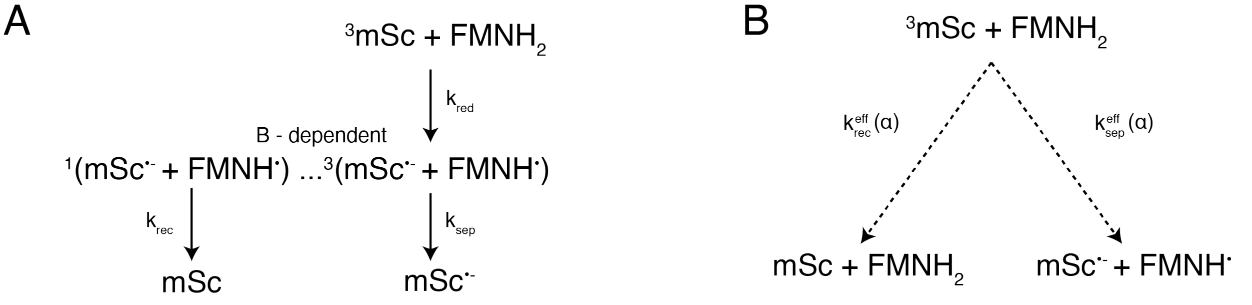


**A)** Triplet mScarlet3 reacts with $FMNH_2$ with rate constant $k_{\text{red}}$ to create a triplet-born SCRP. The singlet SCRP recombines with rate constant $k_{\text{rec}}$ and the triplet SCRP separates with rate $k_{\text{sep.}}$ **B)** The dynamics in **A)** are simplified to a magnetically tuned two-way branch directly from $^3$mSc + $FMNH_2$ to the separation or recombination products.

The rates out of the SCRP are then:

$$k_{\text{sep}} \alpha [\text{SCRP}] = \alpha k_{\text{sep}} \frac{k_{\text{red}}}{\alpha k_{\text{sep}} + (1-\alpha) k_{\text{rec}}} [\text{FMNH}_2][^3\text{mSc}]$$
$$k_{\text{rec}} (1-\alpha) [\text{SCRP}] = (1-\alpha) k_{\text{rec}} \frac{k_{\text{red}}}{\alpha k_{\text{sep}} + (1-\alpha) k_{\text{rec}}} [\text{FMNH}_2][^3\text{mSc}]$$

Therefore, we can make the approximation that the $^3\text{mSc}+\text{FMNH}_2$ reaction goes either directly to the separated radicals, or to the SCRP recombination products, with a magnetic field-dependent branching ratio. Reactions {4}–{6} are replaced by:

$$^3\text{mSc} + \text{FMNH}_2 \xrightarrow{k^{\text{eff}}_{\text{sep}}(\alpha)} \text{mSc}^{\bullet -} + \text{FMNH}^{\bullet} \qquad \{10\}$$

$$^3\text{mSc} + \text{FMNH}_2 \xrightarrow{k^{\text{eff}}_{\text{rec}}(\alpha)} \text{mSc} + \text{FMNH}_2 \qquad \{11\}$$

The effective rate constants are:

$$k^{\text{eff}}_{\text{sep}}(\alpha) \equiv \frac{k_{\text{red}}}{1 + \frac{(1-\alpha) k_{\text{rec}}}{\alpha k_{\text{sep}}}}$$
$$k^{\text{eff}}_{\text{rec}}(\alpha) \equiv \frac{k_{\text{red}}}{1 + \frac{\alpha k_{\text{sep}}}{(1-\alpha) k_{\text{rec}}}}$$

We define $\eta = \frac{k_{\text{rec}}}{k_{\text{sep}}}$. As one would expect, $k^{\text{eff}}_{\text{sep}} + k^{\text{eff}}_{\text{rec}} = k_{\text{red}}$, i.e. the sum of the rates out of the SCRP equals the rate in, at steady state. Furthermore, $k^{\text{eff}}_{\text{rec}} / k^{\text{eff}}_{\text{sep}} = \frac{(1-\alpha)}{\alpha} \eta$, i.e. the branching ratio of the reaction depends on the magnetic-field dependent SCRP S:T ratio, $(1-\alpha)/\alpha$, and also on the ratio of reaction rate constants for the S and T states, $\eta$.

With these simplifications the rate equations are as below. These are simulated in the Supplementary Code:

$$
\begin{aligned}
\frac{d[\mathrm{mSc}]}{dt} &= -k_{\mathrm{ISC}}^{\mathrm{eff}}[\mathrm{mSc}] + \left(k_R^{\mathrm{T}} + k_{\mathrm{rec}}^{\mathrm{eff}}(\alpha)[\mathrm{FMNH_2}]\right)[^3\mathrm{mSc}] + k_{\mathrm{ox}}[\mathrm{FMN}][\mathrm{mSc}^{\bullet-}] && (1)\\
\frac{d[^3\mathrm{mSc}]}{dt} &= k_{\mathrm{ISC}}^{\mathrm{eff}}[\mathrm{mSc}] - k_R^{\mathrm{T}}[^3\mathrm{mSc}] - k_{\mathrm{red}}[\mathrm{FMNH_2}][^3\mathrm{mSc}] && (2)\\
\frac{d[\mathrm{mSc}^{\bullet-}]}{dt} &= k_{\mathrm{sep}}^{\mathrm{eff}}(\alpha)[\mathrm{FMNH_2}][^3\mathrm{mSc}] - k_{\mathrm{ox}}[\mathrm{FMN}][\mathrm{mSc}^{\bullet-}] && (3)\\
\frac{d[\mathrm{FMN}]}{dt} &= -k_{\mathrm{ox}}[\mathrm{FMN}][\mathrm{mSc}^{\bullet-}] + k_{\mathrm{dis}}[\mathrm{FMNH}^{\bullet}]^2 && (4)\\
\frac{d[\mathrm{FMNH_2}]}{dt} &= -k_{\mathrm{sep}}^{\mathrm{eff}}(\alpha)[\mathrm{FMNH_2}][^3\mathrm{mSc}] + k_{\mathrm{dis}}[\mathrm{FMNH}^{\bullet}]^2 && (5)\\
\frac{d[\mathrm{FMNH}^{\bullet}]}{dt} &= [\mathrm{FMNH_2}]k_{\mathrm{sep}}^{\mathrm{eff}}(\alpha)[^3\mathrm{mSc}] - 2k_{\mathrm{dis}}[\mathrm{FMNH}^{\bullet}]^2 + k_{\mathrm{ox}}[\mathrm{FMN}][\mathrm{mSc}^{\bullet-}]. && (6)
\end{aligned}
$$

### 3 Two-state approximation

To determine an analytical form for the dependence of the recovery and quenching rates on [FMN] and $[\mathrm{FMNH_2}]$ (Fig. 6B–E), we also make the approximations $[\mathrm{FMN}] \gg [\mathrm{mSc}^{\mathrm{Tot}}]$ and $[\mathrm{FMNH_2}] \gg [\mathrm{mSc}^{\mathrm{Tot}}]$. These approximations are not universal, but do correspond to the conditions of our *in vitro* experiments, where both flavin concentrations were at least 10-fold greater than $[\mathrm{mSc}^{\mathrm{Tot}}]$. Under these approximations, Eq. 2 becomes:

$$
\frac{d[^3\mathrm{mSc}]}{dt} \approx 0 \approx k_{\mathrm{ISC}}^{\mathrm{eff}}[\mathrm{mSc}] - k_R^{\mathrm{T}}[^3\mathrm{mSc}] - k_{\mathrm{red}}[\mathrm{FMNH_2}][^3\mathrm{mSc}].
$$

Thus $[^3\mathrm{mSc}] = \beta[\mathrm{mSc}]$, with:

$$
\beta \equiv \frac{k_{\mathrm{ISC}}^{\mathrm{eff}}}{k_R^{\mathrm{T}} + k_{\mathrm{red}}[\mathrm{FMNH_2}]}.
$$

Then, we can simplify the dynamics to a two-state model:

$$
\begin{aligned}
\frac{d[\mathrm{mSc}]}{dt} &= -\beta k_{\mathrm{sep}}^{\mathrm{eff}}[\mathrm{FMNH_2}][\mathrm{mSc}] + k_{\mathrm{ox}}[\mathrm{FMN}][\mathrm{mSc}^{\bullet-}]\\
\frac{d[\mathrm{mSc}^{\bullet-}]}{dt} &= \beta k_{\mathrm{sep}}^{\mathrm{eff}}[\mathrm{FMNH_2}][\mathrm{mSc}] - k_{\mathrm{ox}}[\mathrm{FMN}][\mathrm{mSc}^{\bullet-}]
\end{aligned}
$$

The magnetic field acts through its influence on $k_{\mathrm{sep}}^{\mathrm{eff}}$.

The initial quenching rate is the sum of the rates into and out of the mScarlet3 radical anion. Expanding the definitions of $\beta$ and $k_{\mathrm{sep}}^{\mathrm{eff}}$, this becomes:

$$
\begin{aligned}
k_{\mathrm{quench}} &= k_{\mathrm{ox}}[\mathrm{FMN}] + \frac{k_{\mathrm{ISC}}^{\mathrm{eff}}}{(1+\eta\frac{1-\alpha}{\alpha})(1+\frac{k_R^{\mathrm{T}}}{k_{\mathrm{red}}[\mathrm{FMNH_2}]})}\\
&= k_{\mathrm{ox}}[\mathrm{FMN}] + \frac{k_{\mathrm{ISC}}^{\mathrm{eff}}}{(\frac{c_1}{[\mathrm{FMNH_2}]} + c_2)} && (7)
\end{aligned}
$$

where $c_1 \equiv \frac{k_R^{\mathrm{T}}}{k_{\mathrm{red}}}\left(1+\eta\,\frac{1-\alpha}{\alpha}\right) = k_R^{\mathrm{T}}/k_{\mathrm{sep}}^{\mathrm{eff}}(\alpha)$ and $c_2 \equiv 1+\eta\,\frac{1-\alpha}{\alpha} = 1+\frac{k_{\mathrm{rec}}^{\mathrm{eff}}}{k_{\mathrm{sep}}^{\mathrm{eff}}}$.

### 4 Model fitting

#### 4.1 Fitting the intersystem crossing rate and the triplet lifetime

This section pertains to the fitting in Figure 5. In the absence of flavins, there is no charge transfer in our model. Only reactions {1}–{3} occur, and rate equations 1–2 apply. They simplify to:

$$\frac{d[\mathrm{mSc}]}{dt} = -k_{\mathrm{ISC}}^{\mathrm{eff}}[\mathrm{mSc}] + k_R^{\mathrm{T}}[{}^3\mathrm{mSc}]$$
$$\frac{d[{}^3\mathrm{mSc}]}{dt} = +k_{\mathrm{ISC}}^{\mathrm{eff}}[\mathrm{mSc}] - k_R^{\mathrm{T}}[{}^3\mathrm{mSc}].$$

We assume the initial conditions $[\mathrm{mSc}(t=0)] = c_0$ and $[{}^3\mathrm{mSc}(t=0)] = 0$. Then,

$$[\mathrm{mSc}(t)] = \frac{c_0}{k_{\mathrm{ISC}}^{\mathrm{eff}} + k_R^{\mathrm{T}}}\left(k_{\mathrm{ISC}}^{\mathrm{eff}} e^{-(k_{\mathrm{ISC}}^{\mathrm{eff}}+k_R^{\mathrm{T}})t} + k_R^{\mathrm{T}}\right) \tag{8}$$

$$[{}^3\mathrm{mSc}(t)] = c_0 - [\mathrm{mSc}(t)]. \tag{9}$$

We measured the rate of exponential decay $k_{\mathrm{transient}}$ and the steady-state ratio of triplet to singlet as $t \to \infty$, $\left([{}^3\mathrm{mSc}]/[\mathrm{mSc}]\right)_{\mathrm{s.s.}}$ for varying light intensities $I\sigma$ (Fig. 5C).

Fig. 5D shows $k_{\mathrm{transient}} = k_{\mathrm{ISC}}^{\mathrm{eff}} + k_R^{\mathrm{T}} = \frac{k_{\mathrm{ISC}}}{k_{\mathrm{ISC}}+k_R^{\mathrm{S}}} I\sigma + k_R^{\mathrm{T}}$, plotted against $I\sigma$. We fit the slope and intercept to determine $\frac{k_{\mathrm{ISC}}}{k_{\mathrm{ISC}}+k_R^{\mathrm{S}}}$ and $k_R^{\mathrm{T}}$. The best-fit value was $k_R^{\mathrm{T}} = 224\,\mathrm{s}^{-1}$. However, $k_{\mathrm{ISC}}^{\mathrm{eff}} \ll k_R^{\mathrm{T}}$ over the range of light intensities we tested. Therefore, the slope is not well-constrained and we cannot determine $k_{\mathrm{ISC}}$ from this graph.

Fig. 5E shows $\left([{}^3\mathrm{mSc}]/[\mathrm{mSc}]\right)_{\mathrm{s.s.}} = \frac{k_{\mathrm{ISC}}^{\mathrm{eff}}}{k_R^{\mathrm{T}}} = \frac{k_{\mathrm{ISC}}}{k_R^{\mathrm{T}}(k_{\mathrm{ISC}}+k_R^{\mathrm{S}})} I\sigma$ against $I\sigma$. We fit the slope, which provides another measure of $k_{\mathrm{ISC}}$, given $k_R^{\mathrm{T}}$. The best-fit value was $k_{\mathrm{ISC}} = 1.9 \times 10^5\ \mathrm{s}^{-1}$.

#### 4.2 Fitting the rates of oxidation and reduction

This section pertains to the fitting in Figs. 6B–E.

After a period of intense yellow illumination to produce $\mathrm{mSc}^{\bullet-}$, we measured the recovery of mSc at constant [$FMNH_2$], variable [FMN], and magnetic field $B = 0$ (Fig. 4G). In these experiments, we needed to illuminate the sample with some yellow light to monitor the mSc recovery, but we wanted to minimize the dose to avoid driving photochemical production of $\mathrm{mSc}^{\bullet-}$. We compromised by rapidly flickering the yellow light between 0 and 100% at a duty cycle $A = 5\%$.

The rate of [mSc] recovery is similar to the rate of quenching, Equation 7:

$$k_{\text{recovery}} = k_{\text{ox}}[\text{FMN}] + A\frac{k_{\text{ISC}}^{\text{eff}}}{\frac{c_1}{[\text{FMNH}_2]} + c_2},$$

but here the second term is multiplied by $A$, the duty cycle of the light. Since $[\text{FMNH}_2]$ was constant for this set of measurements, we plotted the recovery rates as a function of the FMN concentration. The slope was $k_{\text{ox}} = 3375\ \text{M}^{-1}\,\text{s}^{-1}$.

Then, at constant [FMN], constant light intensity, and variable $[\text{FMNH}_2]$ (Fig. 4H), we measured the initial quenching rates. The magnetic field was $B = 0$.

We plotted $k_{\text{quench}} - k_{\text{ox}}[\text{FMN}] = \frac{k_{\text{ISC}}^{\text{eff}}}{\frac{c_1}{[\text{FMNH}_2]}+c_2}$ against $[\text{FMNH}_2]$. We used the best-fit value of $k_{\text{ox}}$ from above, and we fit the dependence of the quenching rates on $[\text{FMNH}_2]$ to determine $c_1$ and $c_2$. Since there is a degeneracy between $\eta$ and $\alpha$, we assumed that $\alpha(0) = 7/9$. For $k_{\text{ISC}} = 1.9 \times 10^5\ \text{s}^{-1}$ and $I\sigma$[1]$= 1652\ \text{s}^{-1}$ , the best-fit values were $k_{\text{red}} = 5 \times 10^7\ M^{-1}s^{-1}$ and $\eta = 0.84$.

#### 4.3 Fitting the SCRP branching ratios

We then fit $\Delta F/F_{\text{ss}}$ and $\Delta F/F_{t=0}$ against [FMN] and $[\text{FMNH}_2]$ to obtain $\alpha(B_{\text{sat}}) = 0.92$. We included a constant offset of 3% to the fluorescence to account for background. The mScarlet3 radical anion was nonfluorescent under 561 nm excitation; the fluorescence of the quenched product could be entirely attributed to fluorescence from residual mScarlet3. We allowed the light intensity (measured at $\approx 1.5\ \text{W/cm}^2$) to vary in our model; the best-fit value was $I = 1652\ \text{s}^{-1} = 1.7\ \text{W/cm}^2$.

#### 4.4 Fitting the E. coli data

This section pertains to the fitting in Fig. 6G.

Using the same kinetic parameters as for the purified protein, we used the 1) the light intensity dependence of the MFE switching rate for the upstroke $k_{\text{switch}}$ (Fig. 1G) to determine $[\text{FMNH}_2]$, and 2) the steady-state residual fluorescence $F_{ss}/F_{t=0}$ at the lowest light intensity we tested ($I\sigma\ = 145\ s^{-1}$) to determine [FMN]. We adjusted $\eta$ to match the maximal amplitude of the relative MFE, $\Delta F/F_{ss}$.
First, we note that the MFE switching rate[2]:

$$\begin{aligned} k_{\text{switch}} = k_{\text{quench}} &= k_{\text{ox}}[\text{FMN}] + \frac{k_{\text{ISC}}^{\text{eff}}}{(\frac{c_1}{[\text{FMNH}_2]} + c_2)} \\ &= k_{\text{ox}}[\text{FMN}] + \frac{I\sigma k_{\text{ISC}}}{(k_R^S + k_{\text{ISC}})(\frac{c_1}{[\text{FMNH}_2]} + c_2)}. \end{aligned}$$

We plot $k_{\text{switch}}$ for the upstroke against $I\sigma$. The best-fit slope was $m = 3.13 \times 10^{-4}$. Using the rate constants from the model fit for the purified protein, we obtained $[\text{FMNH}_2] = 6\ \mu\text{M}$. However, the error on the intercept

[1]The molar extinction coefficient of mScarlet3 at 561 nm is $\epsilon_{561} = 89,440\ M^{-1}\text{cm}^{-1}$ which corresponds to an absorption cross section of $\sigma_{561} = \frac{\ln(10)\times 10^3}{N_A}\epsilon_{561} = 3.46\ \times 10^{-16}\ \text{cm}^2$.
[2]For a discussion of the switching rate, see Section 5.2.

was too large to determine [FMN].

Then, we note that the quantity:

$$
\begin{aligned}
F_{ss}/(F_{t=0} - F_{\mathrm{ss}}) &= \left(\frac{[\mathrm{mSc}]}{[\mathrm{mSc}^{\bullet-}]}\right)_{\mathrm{ss}} \\
&= \frac{k_{\mathrm{ox}}[\mathrm{FMN}](\frac{c_1}{[\mathrm{FMNH_2}]} + c_2)}{k^{\mathrm{eff}}_{\mathrm{ISC}}} \\
&= \frac{k_{\mathrm{ox}}[\mathrm{FMN}]}{I\sigma\ m}
\end{aligned}
$$

where $m$ is the slope from the fit previously. Rearranging, $[\mathrm{FMN}] = \frac{I\sigma\ m}{k_{\mathrm{ox}}}\frac{F_{\mathrm{ss}}}{F_{\mathrm{t=0}} - F_{\mathrm{ss}}}$. At $I\sigma = 144\ \mathrm{s}^{-1}$, $F_{ss}/(F_{t=0} - F_{\mathrm{ss}})$ $= 0.03$. We choose to use the lowest light intensity to reduce the impact of photobleaching on the measurement of $F_{\mathrm{ss}}$. We obtained [FMN] = 0.4 $\mu$M.

### 5 Maximizing the MFE amplitude and kinetics

#### 5.1 MFE amplitude

We seek a formula for the absolute and relative MFE amplitudes, $\Delta F/F_{\mathrm{t=0}}$ and $\Delta F/F_{\mathrm{ss}}$, respectively. Using the approximation from Section 3, we write the fractional steady-state residual fluorescence $x \equiv F_{\mathrm{ss}}(\alpha)/F_{\mathrm{t=0}}$ as:

$$
\begin{aligned}
x &= \frac{k_{\mathrm{ox}}[\mathrm{FMN}]}{k_{\mathrm{quench}}} \\
&= \frac{k_{\mathrm{ox}}[\mathrm{FMN}]}{k_{\mathrm{ox}}[\mathrm{FMN}] + \frac{k^{\mathrm{eff}}_{\mathrm{ISC}} \cdot k_{\mathrm{red}}[\mathrm{FMNH_2}]}{\left(1+\eta\frac{1-\alpha}{\alpha}\right)\left(k_{\mathrm{red}}[\mathrm{FMNH_2}] + k^{\mathrm{T}}_{R}\right)}} \\
&= \frac{k_R}{k_R + \frac{k_F}{(1+\eta\,\frac{1-\alpha}{\alpha})}}
\end{aligned}
$$

where we have defined $k_R$ as the rate out of the mScarlet3 radical anion, and $k_F$ as the non-$\alpha$-dependent portion of the rate into the radical anion. We take the derivative with respect to $\alpha$:

$$
\begin{aligned}
\frac{dx}{d\alpha} &= \frac{-k_R k_F}{\left(k_R + \frac{k_F}{(1+\eta\,\frac{1-\alpha}{\alpha})}\right)^2} \cdot \frac{1}{(1+\eta\,\frac{1-\alpha}{\alpha})} \cdot \frac{\eta}{\alpha(\alpha + \eta - \alpha\eta)} \\
&= -x(1-x)\frac{\eta}{\alpha(\alpha+\eta-\alpha\eta)} \\
&= -x(1-x)\frac{\eta}{\alpha^2(1+\eta\frac{1-\alpha}{\alpha})}.
\end{aligned}
$$

Therefore, in the limit where $\Delta\alpha \equiv \alpha(B_{\text{sat}}) - \alpha(0)$ is small,

$$\Delta F/F_{t=0} = \frac{\eta\Delta\alpha}{\alpha^2\left(1+\eta\frac{1-\alpha}{\alpha}\right)}x(1-x) \tag{10}$$

$$\Delta F/F_{\text{ss}} = \frac{\eta\Delta\alpha}{\alpha^2\left(1+\eta\frac{1-\alpha}{\alpha}\right)}(1-x). \tag{11}$$

#### 5.2 Fluorescence quenching and MFE switching kinetics

Upon a change in the magnetic field (in the model, a change in $\alpha$), the fluorescence reaches a new steady-state, $F_{\text{ss}}$, depending on the new rate into the radical anion. $k_{\text{switch}}$ is the rate of $\text{mSc}^{\bullet -}$ reaching steady-state. At low light intensity ($I\sigma \lesssim 10^6$ s$^{-1}$), the quenching rate $k_{\text{quench}}$ is the same:

$$k_{\text{switch (upstroke)}} = k_{\text{quench}}(\alpha(0)) \tag{12}$$

$$k_{\text{switch (downstroke)}} = k_{\text{quench}}(\alpha(B_{\text{sat}})). \tag{13}$$

$k_{\text{quench}}$ and thus $k_{\text{switch}}$ are linearly dependent on [FMN] and $I\sigma$, and also depend on [$FMNH_2$] up to $c_1$ (Equation 7) which is $\approx 5.5$ $\mu$M. Therefore, to increase the MFE switching rate, one should increase the light intensity and flavin concentrations. We note that the choice of which parameter to increase affects the residual fluorescence $F_{\text{ss}}/F_{\text{t=0}}$ (and subsequently, the MFE amplitude) differently. Increasing [FMN] will increase $F_{\text{ss}}/F_{\text{t=0}}$, while increasing the light intensity or [$FMNH_2$] will decrease it.

At high light intensities and high frame rate, the definition of quenching changes. Upon illumination, a substantial portion of the initial fluorescence drop is attributed to populating photocycle intermediates — the mScarlet3 triplet state $^3$mSc (at $I\sigma \gtrsim 10^6$ s$^{-1}$)[3] and the excited state, mSc* (at $I\sigma \gtrsim 10^8$ s$^{-1}$).

Our simulations of the switching as a function of light intensity in Fig. S13 explicitly track the excited state, [mSc*] and fit the upstroke switching rate. The range of $I\sigma$ over which Eqs. 12–13 will hold generally depends on [$FMNH_2$], since the rate out of the mScarlet3 triplet state is [$FMNH_2$]-dependent. We have stated the approximate ranges for [FMN]= 30 $\mu$M and [$FMNH_2$] = 10 $\mu$M.

[3] The model predicts that if $I\sigma = 10^6$ s$^{-1}$, [FMN] = 30 µM, [$FMNH_2$] = 10 µM, the dynamics are significantly different from what we observed in experiments. During the initial drop in fluorescence, [$^3$mSc] builds up. [$^3$mSc]/[$\text{mSc}^{\text{Tot}}$] peaks at 40% and then $^3$mSc slowly converts to $\text{mSc}^{\bullet -}$. The timescale of this is slightly faster comparable to the MFE switching rate at this light intensity, thus the quenching is described by a multiexponential. Subsequent switching events after a long time are accompanied by smaller variations in $^3$mSc. The approximation in Section 3 no longer holds.
